# “Axonal Length Determines distinct homeostatic phenotypes in human iPSC derived motor neurons on a bioengineered platform”

**DOI:** 10.1101/2021.08.30.458271

**Authors:** Cathleen Hagemann, Carmen Moreno Gonzalez, Ludovica Guetta, Giulia Tyzack, Ciro Chiappini, Andrea Legati, Rickie Patani, Andrea Serio

**Author notes:** corresponding author ( /).

## Abstract

Stem cell-based experimental platforms for neuroscience can effectively model key mechanistic aspects of human development and disease. However, conventional culture systems often overlook the engineering constraints that cells face *in vivo*. This is particularly relevant for neurons covering long range connections such as spinal motor neurons (MNs). The axons of these neurons extend up to 1m in length and require a complex interplay of mechanisms to maintain cellular homeostasis. It follows that shorter axons in conventional cultures may not faithfully capture important aspects of their longer counterparts. Here we directly address this issue by establishing a bioengineered platform to assemble arrays of human axons ranging from micrometers to centimeters, permitting systematic investigation of the effects of length on human axonal biology for the first time. With this approach, we reveal a link between length and metabolism in human MNs *in vitro*, where axons above a “threshold” size induce specific molecular adaptations in cytoskeleton composition, functional properties, local translation and mitochondrial homeostasis. Our findings specifically demonstrate the existence of a length-dependent mechanism that switches homeostatic processes within human MNs in order to sustain long axons. Our findings have critical implications for *in vitro* modelling of several neurodegenerative disorders and reinforce the importance of modelling cell shape and biophysical constraints with fidelity and precision *in vitro*.

## 1. Introduction

Shape and function are two intimately linked properties in all living cells: the need to perform a certain function configures each cell to a specific and optimal shape, which in turn also influences cellular homeostasis^1^. In cells with polarized or complex shapes several key functions need to be performed remotely from the soma, which creates challenges that sometimes cannot be overcome by soley relying on diffusion of metabolites and components. While in principle cell shape is difficult to study systematically *in vitro* for a specific cell type, recently a combination of stem cell technology advancements and bioengineering techniques have substantially galvanized this field of investigation. Thanks to reliable protocols of differentiation and the advent of cellular reprogramming we can generate most cell populations, which has revolutionized discovery science in human biology. Neuroscience in particular has benefitted substantially from the possibility of creating reproducible *in vitro* models to both better understand neural cell biology and model diseases.

To complement this revolution, bioengineering tools have been developed and made accessible to direct cell growth, alter cell shape and their relative position, thus enabling the development of on-chip devices based on microfluidics systems or microfabricated topography.

Neurons have a highly polarized architecture^2^ and are also amongst the largest known cells^3^ with their main sites of activity and high energy demand in the periphery (e.g. synapses). Spinal **Motor Neurons (MNs)** in particular can have axons of up to 1 m in length in humans^4^, which poses significant challenges to the maintenance their homeostatic processes and function. For example, metabolic processes and mitochondrial activity are needed to produce ATP, which in turn is used to maintain axonal transport, membrane potential and all other functions; but across long distances simple diffusion and motor-based transport may cause too slow a response to sudden changes in metabolic demands. Therefore, neurons have evolved adaptive mechanisms to maintain these and other essential processes in the axon and terminals^3^, from localized control of mitochondrial homeostasis - or mitostasis^5^- and dynamics^6^, axonal RNA targeting, local protein translation (LPT)^7,8^ and alterations to the axonal cytoskeleton^9^.

Interestingly, axonal mitochondrial and metabolic breakdown are recognized early phenotypes of neurodegeneration and peripheral neuropathies (amyotrophic lateral sclerosis-ALS^10,11^ or Charcot-Marie-Tooth, CMT^12^), as well as in other pathologies (e.g. in long nerve bundles in experimental diabetic neuropathy^13^). Moreover, a general decline in metabolic control, mitostasis and transport in the axon is also a common feature in normal ageing^14^.

Accurately modelling MNs *in vitro* is crucial given that they are the target of several incurable neurodegenerative diseases, (e.g. ALS, CMTs and others). We and others have over the years developed several stem cell *in vitro* platforms for MNs combining stem cells, bioengineering, and imaging^10,15–18^ to faithfully recapitulate their biology and mechanisms of disease in ALS. However, these *in vitro* models fail to recapitulate the length of motor axons, which is in most cases the used culture systems generate cells with axons ranging from 1-200um^19^ to at most 1 millimeter ^20,21^ or do not provide a controlled environment for measuring axonal length, lacking the modelling of potential biophysical constraints which long axons provide *in vivo*.

Given this gap between these cultures and their *in vivo* counterparts, whether physical axonal length is a salient property to model for MNs - and by extensions other neurons-*in vitro* is a key question that remains unresolved. Addressing this issue may have important implications for *in vitro* experimental studies examining basic cellular process or disease biology.

However, at present no systematic studies have been conducted to explore this problem, largely due to technological limitations. While several axonal chambers and microfluidic devices have been developed to study soma/axonal differences, they tend to be either enclosed system that can only produce ordered arrays between open chambers, with no control over the actual length of the axons and cannot easily be adapted to answer this question. We therefore sought to create an ideal system to study axonal length systematically with human neurons *in vitro* and investigate length-dependent alterations in axonal homeostatic processes, where: i) the only factor changing is axonal length and one can control it reliably; ii) the whole axon is accessible.

In this study we report the establishment of a bioengineered platform and its use to conduct the first systematic study on the molecular effects of axonal length on internal homeostatic dynamics within the axoplasm. We reveal that length determines specific alterations to metabolic, functional, and structural properties of motor neurons *in vitro* and therefore demonstrate that axonal length is a salient characteristic to model in both basic and applied studies of neurons.

## 2. Results

### 2.1. Combining microtopography and neural aggregate cultures to optimize an axonal elongation platform

To generate a platform that allows us to systematically investigate the effects of axonal length on neuronal homeostasis, we first sought to optimize a protocol to grow human neurons on bioengineered, through soft-lithography generated, substrates, which provide guidance of axonal elongation and directionality^20,21^.

We have previously established a directed differentiation system to generate functional spinal MNs from iPSC populations^10^ and several protocols and pipelines to assess different ALS phenotypes - including axonal specific ones-within these cultures^18,22^ showing that the stem cell-based systems recapitulate several key aspects of the disease. We have therefore started this investigation by employing our established differentiation protocols to obtain homogenous populations of spinal MN progenitors, which we cultured to confluency and then transitioned from a 2D system to an organoid-like 3D aggregated culture, by allowing for an extra 48h expansion past the usual split point (c.a. 80% confluency). This generates tightly packed groups of progenitors within the culture, which are then lifted and separated from the surrounding single cells using PBS-EDTA and transferred to a suspension culture on an orbital shaker enhancing the aggregate formation and preventing cell attachment (**Figure 1A**). The generated 3D MN aggregates have a homogeneous size (approx. 200μm in diameter with regular spherical shape and consistent in size, **Figure S1**) and can be immediately differentiated in suspension, by withdrawing FGF2 for 48h, and then plated on cell culture substrates to obtain a dense aggregates of cell bodies that quickly grow axons out of the 3D structure.

**Figure 1:**
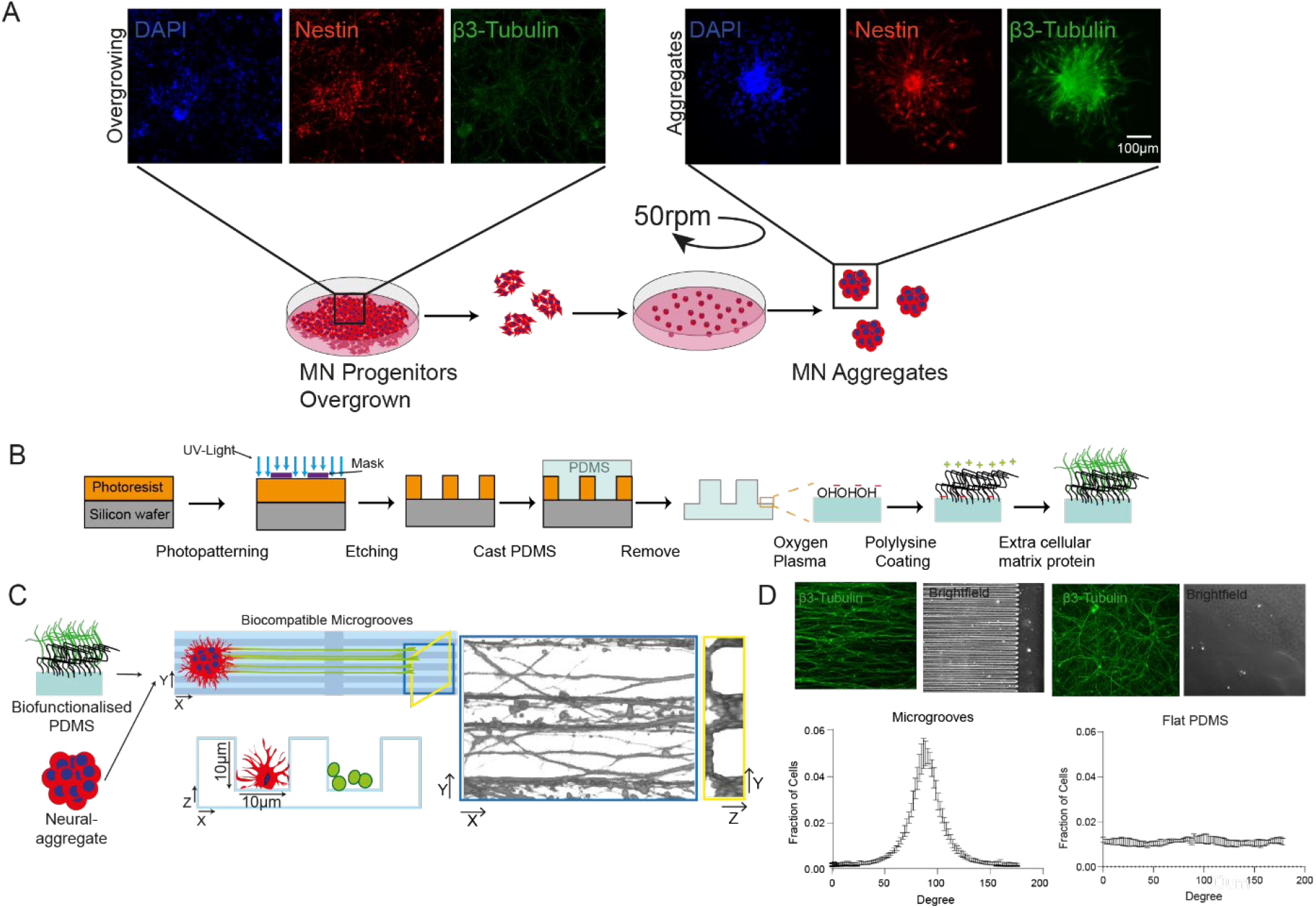
Overview of the long axon motor (LAM) array protocol and cell alignment. **(A)** hIPSC derived motor neuron progenitors are cultured as previously described. Cells are then overgrown for 2 days past their confluency point and used to generate neural aggregates by suspension culture on an orbital shaker. Immunofluorescence panels show representative images of the MN progenitors expressing relevant developmental markers. NESTIN for progenitors and β-3 TUBULIN for differentiated neurons and DAPI for nuclei at each stage. **(B)** Schematic overview of the protocol applied to generate micropatterned surfaces using soft-lithography and biofunctionalization of PDMS. For generation of a patterned surface on a silicon wafer SU8 is used and photopatterned using a striped mask and UV light. An imprint of this pattern is created using the biocompatible material PDMS. PDMS is oxygen plasma treated and poly-D-lysine linked and then coated with an extra cellular matrix protein as laminin, allowing cell attachment. **(C)** Assembly of long axon motor arrays using neural aggregates and biofunctionalized micro patterned surfaces. The surface has a microgroove pattern (10umx10um, w × h) interspersed with a 250um wide unpatterned area (plateau). Neural aggregates were pre-differentiated in suspension and then placed with a pipette onto the biofunctionalized substrate. Upon differentiation, axons extend from the aggregates along the grooves. Right panel show a 3D confocal reconstruction in x-y direction (blue) and y-z (yellow) of axons cultured on the substrate using β-3 tubulin. **(D)** Comparison of cell directionality on patterned and flat PDMS, showing alignment of cells to the given topography. No alignment is achieved on a flat substrate and representative fluorescent images of the cells on the different substrate are displayed in the top panel, which were used for analysis.

We have combined these 3D neural aggregates with 10µm × 10µm (width × height) microgrooves imprinted in PDMS using soft-lithography^23^. Additionally, we inserted every 5 mm a 250µm wide flat plateau to control for axonal elongation using brightfield microscopy. The micropatterned PDMS is then biofunctionalized using a combination of poly-D-lysine and laminin (**Figure 1B-C**) after oxygen plasma treatment, to generate ordered arrays of axons. We have focused first on optimizing the fabrication, functionalization, and culture time of the aggregates on the grooves to maximize the effect of the microtopography on axonal length (**Figure 1 D & Figure S2**). This optimization is essential to create a platform that systematically directs axonal growth, maintaining axonal length as the sole biophysical variable. As the stiffness of PDMS can be adjusted by varying the ratio of curing agent to monomer, we tested different compositions and different biofunctionalization molecules to optimize our devices for axonal outgrowth and found that PDMS 1:10 is the most suitable substrate, yielding the longest neurons within a given timeframe, and with the addition of microgrooves it provides axonal guidance using topographical features (**Figure S3**), which allows to obtain ordered arrays of axons and simplify subsequent imaging approaches to analyze specific points across the length. This system, which we have named Long Axon for Motor neurons (LAM) arrays, allows to obtain significantly longer and oriented axons than any of the unoptimized cultures.

Once optimized, the LAM arrays allow us to effectively direct axonal elongation and maintaining separation of cell bodies, dendrites, and axons. Due to the properties of the 3D aggregate culture, dendrites tend to stay close to the cell bodies and remain significantly shorter compared to axons, which instead interact with the microgrooves and quickly outgrow the other neurites (**Figure 2A-B**). This effectively creates a simple and reproducible system to isolate axons from the soma for investigation without requiring physical barriers. By varying the time in culture for the LAM arrays we can obtain cultures with identical starting conditions that have as the only variable the scale of axonal length, which can be effectively tuned from several hundreds of µm (hence representative of conventional cultures), to several mm, up to and exceeding 1 cm within 3-4 weeks of culture, while still allowing to investigate single axons within the culture (**Figure 2D-E**).

**Figure 2:**
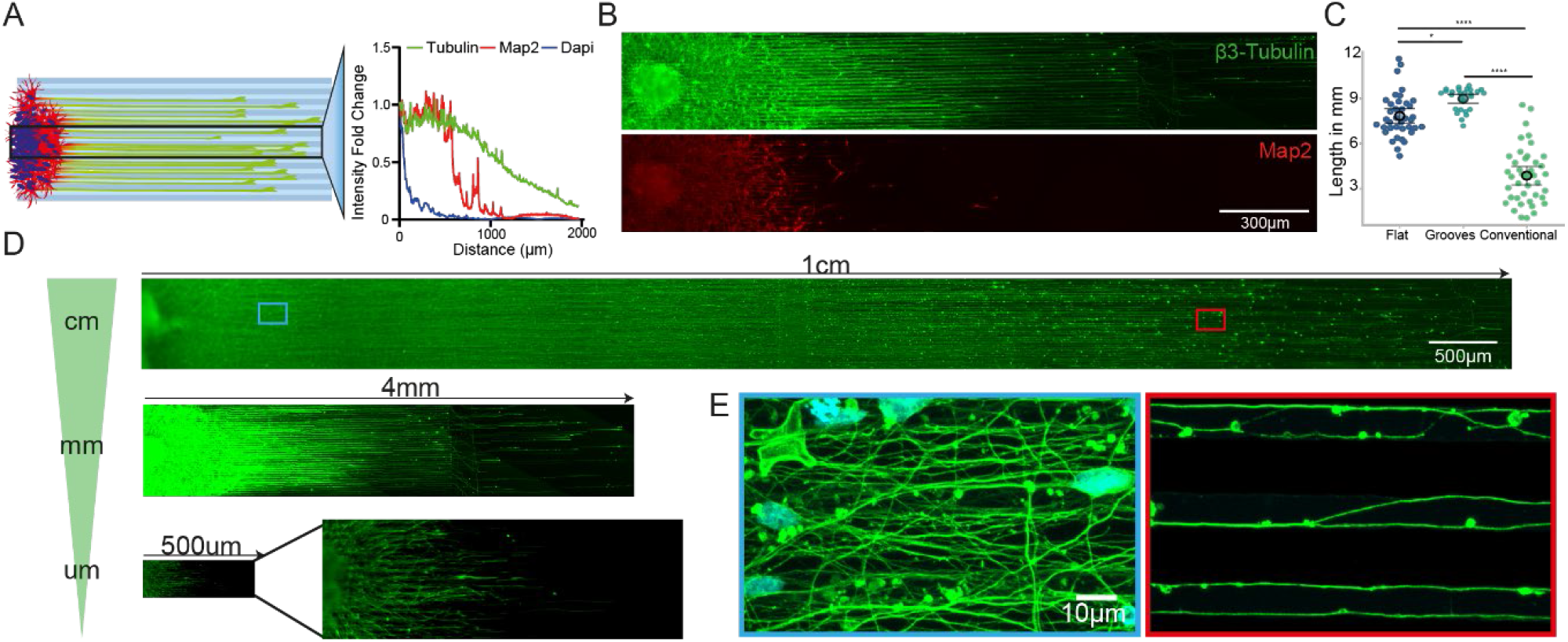
LAM arrays allow to obtain different scaled cultures of motor neuron axons. **(A-B)** Schematic representation of line profile measurements performed on LAM arrays stained for axonal β-3 TUBULIN, dendritic (MAP2) and nuclear (DAPI) markers 14 days after seeding (A). The graph on the right shows a representative line profile plot across β-3 TUBULIN positive axons, showing that dendrites are solely located in close proximity to the aggregate whereas axons extend throughout the microgrooves with limited cell migration. (B) Representative fluorescent images of axonal and dendritic staining of LAM arrays with long motor axon cultures 14d in differentiation **(C)** Culturing of axons on microgrooves improves overall length and homogeneity compared to flat PDMS or conventional culturing on tissue culture plastics after 3 weeks in culture. Measurement of axonal length in mm on different substrates using β-3 TUBULIN staining or single cell transfections with GFP (conventional) across single axons. (Kruskal-Wallis test *p=0.0182, ****p<0.001, Mean+- 95%CI calculated on 1 independent experimental block with 1 cell lines each with 3 technical replicates). **(D-E)** Using LAM arrays allows culture of axons in the cm-range after 21d in differentiation or mm-range after 14d in differentiation. µm-ranged motor axons are achieved after 1d in differentiation. Representative fluorescent images of the different axonal length achieved on LAM array cultures are immunocytochemistry stained for β-3 TUBULIN and are presented with the same pixel size (D). Representative confocal images of β-3 TUBULIN positive long motor axons on bioengineered substrates close to the aggregate (blue) and distal regions (red) projecting along the topography (E).

### 2.2. LAM array analysis shows structural and functional alteration that arise past specific “threshold” length in vitro

Using the LAM platform, we sought first to understand whether structural components within the axons would change in distribution or composition at different axonal lengths. The most fundamental structural component for axons is the cytoskeleton, and we focused in this instance on neurofilaments (NFs) as they are highly abundant in axons. Neurofilaments are arranged in a tripartite structure, consisting of differently sized subunits which are named after their respective weight, light (NFL), medium (NFM) and heavy (NFH) chains. Their composition and abundance has been shown to change between species and tissues but also along single axons *in vivo* but less is known about their relative changes within human motor axons and the effect of their abundance on axonal length *in vitro*. We therefore sought to analyze the distribution and ratio between light (NFL) and heavy chain (NFH) along human motor axons, which were grown to 1 cm in length on our LAM devices. Fluorescent intensities of the respective proteins were normalized to the β-3 TUBULIN signal accounting for axonal area and number. Interestingly, we observed that while NFL remains stable along the axons, NFH has a dual mode distribution, decreasing rapidly from the µm to mm scale and stabilizing around 4-5 mm to the level of NFL until the axonal terminal is reached (**Figure 3A**). To investigate the composition, we further calculate the ratio of the two neurofilament subunits comparing proximal, medial, and distal axonal compartments, which show a continuous change with NFL becoming the predominant subunit. This confirmed the single subunit change and the correlation with the length of the axon (**Figure 3B-D**). We then wanted to check whether other structural and functional components within long axons display a similar length-dependent alteration in their distribution. The endoplasmic reticulum (ER) is a key component of the axoplasm it consists of a physically connected continuous membrane bound organelle that spans the entirety of the axon contacting other organelles and the cytoskeleton, and it serves numerous specialized roles as protein export, calcium storage or lipid synthesis^24,25^. Calreticulin is a major component of the ER and, similarly to what we observed for NFH, is significantly affected by axonal length. In axons reaching 1 cm calreticulin is showing marked changes in distribution compared to much shorter ones when considering whole axons **(Figure 3E)** and in a compartment specific manner **(Figure 3F-G)**.

**Figure 3:**
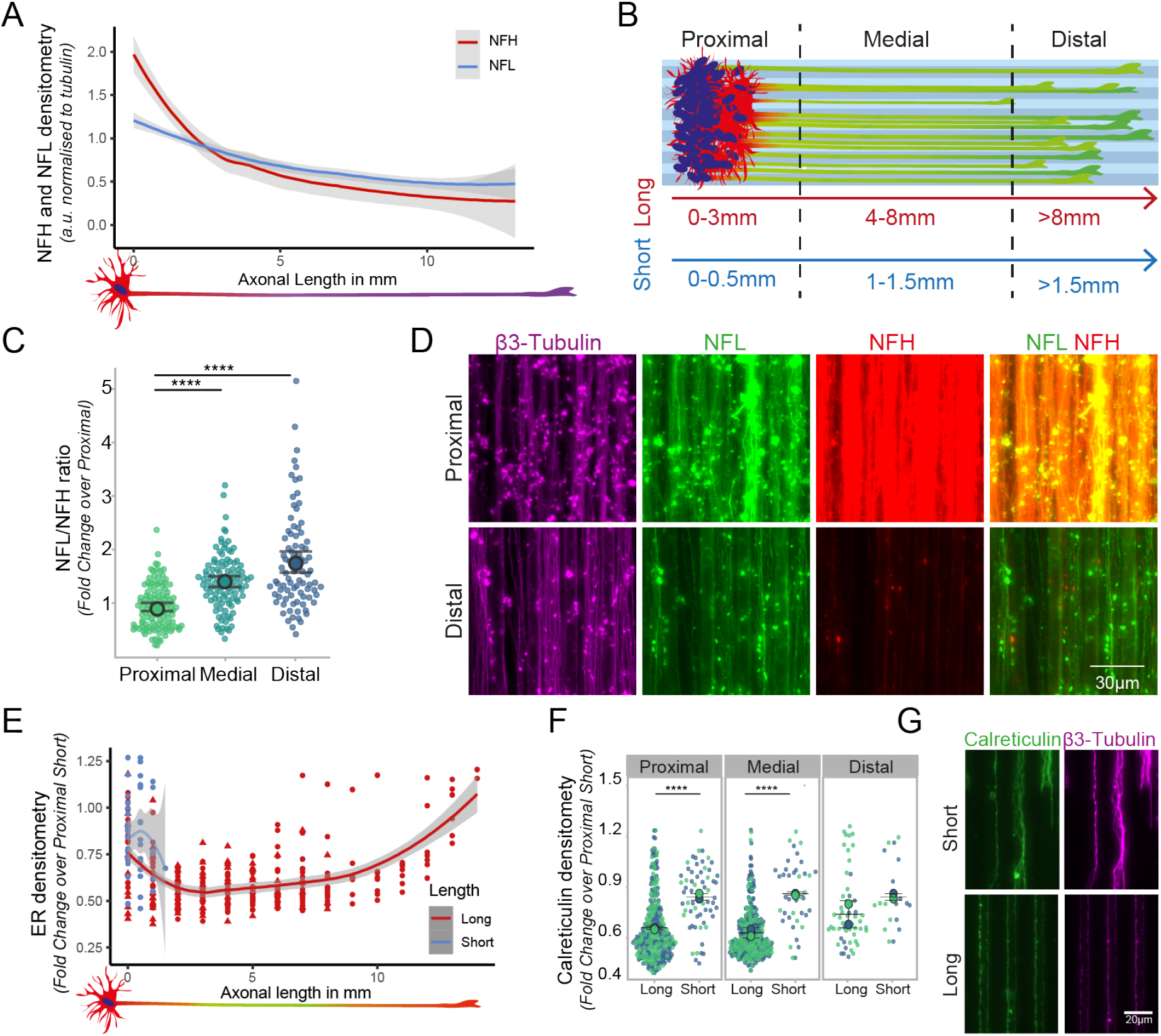
LAM array cultures allow systematic analysis of length dependent changes in internal composition. **(A)**Long axons cultured on LAM arrays show different distribution of the neurofilament subunits, light (NFL) and heavy (NFH) chain. Line graph shows NFL (blue) and NFH (red) fluorescent intensities normalized to β-3 TUBULIN. **(B)** To systematically compare compartment specific effects, axon arrays of different lengths are separated into proximal, medial, and distal segments, relative to their total axonal length on LAM arrays (color code). **(C)** The ratio between the neurofilament subunits is gradually changing in a compartment specific manner. 1 independent experimental block with 2 cell lines each with 3 technical replicates and 2 independent experimental blocks with 1 cell line each with 3 technical replicates. (Kruskal-Wallis test with Dunn’s multiple comparison ****p<0.0001.) **(D)**. Fluorescent images of stained compartments for different neurofilament subunits and β-3 tubulin. **(E-G)** Quantification of CALRETICULIN staining density normalized to β-3 tubulin, indicates length dependent alterations in ER-axonal presence and distribution compared to short LAM arrays, both at the whole axon level (E), and within specific compartments (F) expressed as fold change compared to the proximal segment of short axons. The different colors represent here the experimental blocks. Representative fluorescent images of calreticulin and β-3 tubulin in short and long axons cultured on bioengineered substrates (G). Analysis conducted on 2 independent experimental blocks with 2 cell lines each with 3 technical replicates. (Mann-Whitney, ****p<0.0001, Mean +- S.D.).

As one of the axonal ER main roles is to regulate calcium (Ca^2+^) storage and signaling, we next sought to investigate whether long motor axons *in vitro* display a length- or compartment-dependent alteration in their intracellular Ca2+ dynamics, which are also a fundamental part of neuronal activity. Using a calcium sensitive Fluo4-AM dye we stained 1cm long motor axons grown on our LAM arrays and performed a live imaging analysis of neuronal firing within our cultures, comparing the different axonal compartments. We acquired images every 100 ms for a total of 100 frames, focusing on the axons using SiR-Tubulin, a live dye that is staining the microtubules, and performed peak detection and frequency analysis using a publicly available Matlab code called ‘PeakCaller’ as previously described^24^. Comparing the peak intensity and distribution, a marked difference between proximal and distal axons becomes immediately evident, with the latter exhibiting decreased spike frequency and height. **(Figure 4A-C)**. Further analysis on the number of calcium transients detected within each axonal compartment and the mean height of the detected spikes revealed a progressive decrease in both measurements while moving from the proximal to the distal compartment of the axons (**Figure 4D-E)**.

**Figure 4:**
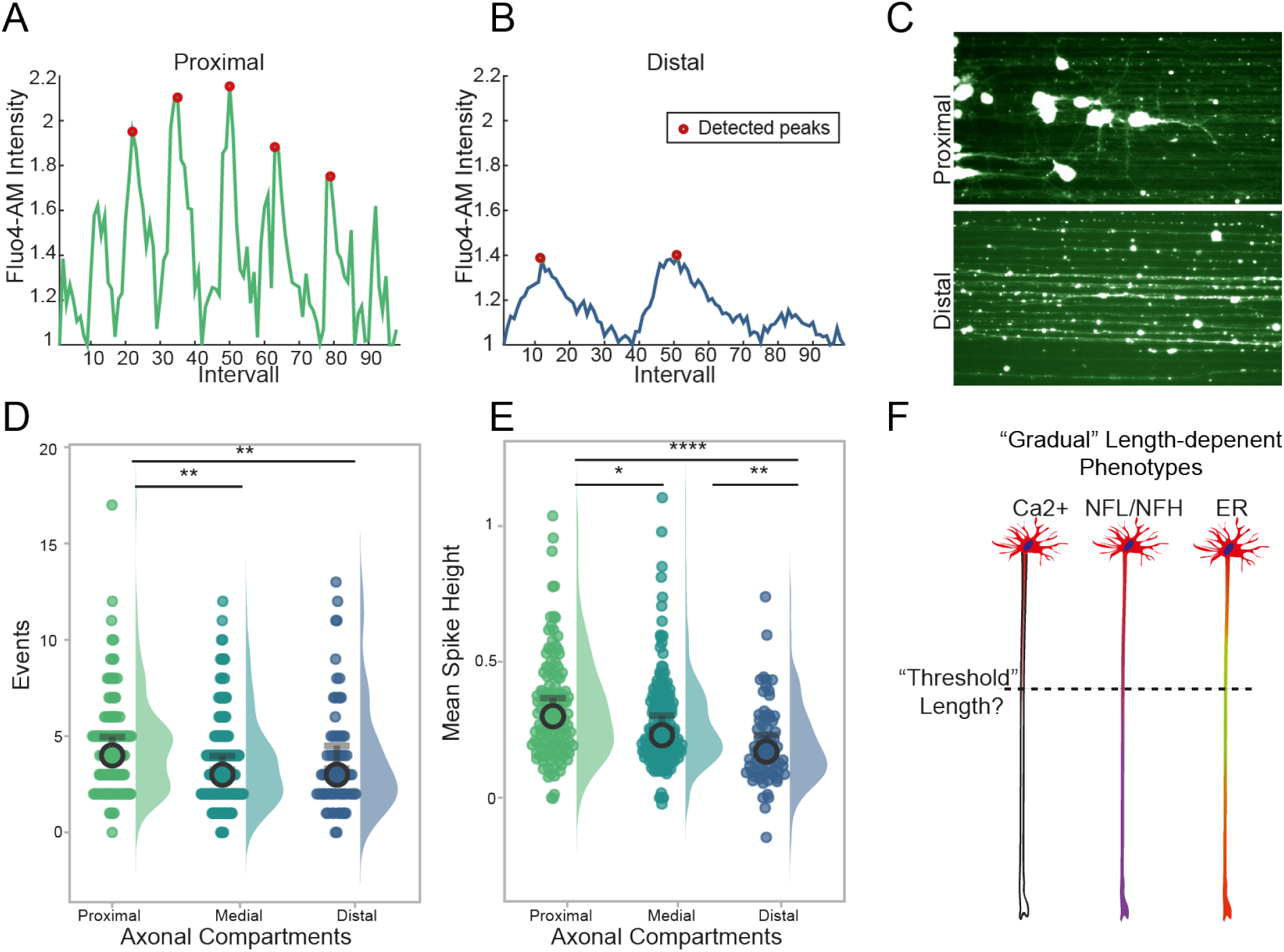
Long human motor axons on the LAM arrays are functionally active and show gradual compartment-dependent changes in calcium signaling across their length. **(A-C)** Fluorescent imaging of calcium using Fluo4am loaded long axon motor arrays is performed on LAM arrays using long motor axons. Representative traces of proximal and distal calcium waves with annotation of detected peaks (redpoint), which have been quantified using a publicly available Matlab code called ‘PeakCaller’ (A-B) Representative images of living long axons on LAM arrays stained with Fluo4AM using “thalium” LUT (C). **(D-E)** Fluo4Am loaded long axons are imaged every 100 ms for 100frames. Quantification of calcium events (D), which are indicated by increased fluorescent signals, are then detected as spikes as above described, (Kruskal Wallis with Dunn’s multiple comparison, **proximal vs distal p=0.0029, proximal vs medial p=0.0014) and mean spike height (E) (Kruskal Wallis test with Dunn’s multiple comparison. *p= 0.0133, **p=0.0012, **** p< 0.0001 Mean +- 95%CI, 1 independent experimental block with 1 cell lines each with 3 technical replicates). **(F)** Threshold hypothesis based on gradual length dependent changes found in fundamental structures of long axons.

Interestingly, our results thus far consistently indicate that it is possible within human motor axons grown to 1 cm *in vitro* to detect gradual length-dependent alterations, that while not indicative of cellular stress or pathological conditions within the neurons, all point to a fundamental change in behavior that becomes evident between the medial and distal portion of the axons, approx. 4-5 mm from the terminals. These observations were biologically consistent, as ER, calcium and cytoskeletal structures are linked^25,26^ and they raise the intriguing possibility that above a certain “threshold” length **(Figure 4F)**, isolated axons need to adapt their homeostatic mechanisms, to accommodate a degree of molecular bottlenecking determined by the lengthening of transport times for organelles, RNA, proteins, and nutrient^2,27^. The fact that these changes appear more evident within the distal axons could indicate compensatory mechanisms that are out in place to maintain neuronal activity at the most distant portion of the cell from the soma.

### 2.3 Length-dependent alterations in axonal mitochondria distribution and shuttling

To further investigate whether axonal length is a major determinant for the proposed biophysical constraint, which might influence homeostatic mechanisms, we sought to investigate two fundamental metabolic processes within axons: mitochondria dynamics and local protein translation. Based on the analysis performed above, we defined 2 distinct groups for our following investigation: neurons with axons below the observed “threshold length” of 4-5mm – which we have classified as “short” (2-3mm in length)- and neurons with axons reaching 8-10 mm and above - which we have classified as “long”.

Mitochondria are essential to the survival and function of any cell, but in particular in neurons they are responsible for generating 90% of ATP^28^ and play a crucial role in providing energy for axonal transport and synaptic function ^29^. First, we sought to investigate the mitochondrial content of short and long axons (as defined above) cultured on LAM arrays, using TOMM20 (Transporter of the Outer Mitochondrial Membrane) as a mitochondrial marker and quantify the axonal area occupied by mitochondria within axons, which was measured by using the area of β-3 tubulin positive structures. Calculating the overall density of mitochondria, expressed as fold change over the proximal compartment of short axons, shows that these organelles are homogenously distributed across long axons but present a decreasing trend in short axons **(Figure 5A)**. Separating the axons into different compartments (proximal, medial, distal) further reveals a significant increase of mitochondrial content within short axons when comparing medial and distal compartments **(Figure 5B)** which is not mirrored in long ones. This length-dependent increase of axonal mitochondria is both significant and with several potential implications for the field, so we wanted to further investigate whether the observed changes are indeed solely dependent on the length of the axon or also influenced by the maturation state and time in culture (which unavoidably differs between long and short axons on our LAM platform, see paragraph 2.1 and Figure 1 for details). Hence, we engineered a specific control experiment where we cultured for 3 weeks axons either on LAM arrays, flat PDMS functionalized as the LAM arrays and in conventional culture plates. We then transfected MNs with GFP to be able to measure axonal length in the conventional culture where the lack of directionality of the axons makes it impossible to assess accurately axonal length at the population level. After the same time in culture across the different conditions, we measured the mitochondrial content as described above and the average axonal length. This experiment firstly confirmed that LAM arrays allow to obtain significantly longer neurons *in vitro* compared to conventional culture systems, and further validated the relationship between axonal length and mitochondrial content, with the longest axons having an increased mitochondrial content independently of the time in culture (**Figure S4**). This characterization of mitochondrial content provides key insights into the effects of axonal length but does not fully capture the complex homeostasis of mitochondria in a cell for maintenance of a balanced system. Mitochondrial transport in the axon is used for redistribution of mitochondria in the cells as a respond to local demands, like neural activity, and other various processes^1^, and - more importantly, it has been shown to change in response to both maturation and compartment. ^29,30^. To track these movements, we stably transfected MN progenitors with a fluorescent photo switchable protein (Dendra2) fused to the Cox8a subunit of mitochondria (Mito)^29^ using a transposon-based PiggyBac vector^30^ (see 5.Experimental Methods for details). The transfected cells were cultured and seeded on LAM arrays to generate short and long axons as described above. Fluorescently tagged mitochondria were live imaged on the arrays every 10s for 2 minutes after reaching the desired length using imaging medium (see 5. Experimental Methods for details) **(Supplementary Video 1-2)**. The resulting time-lapse images were converted into kymographs **(Figure 5C)**, to investigate the shuttling of mitochondria along the axons. We first focused on determining the proportion of stationary vs. motile mitochondria, which correlates with axonal maturity and with overall local ATP demands in the axons^30^. We found that significantly more mitochondria are stationary in long axons with an average ∼ 92%, which is in line with the above-mentioned maturation profile in axons. We then focused on the fraction of moving mitochondria to investigate their directionality. Mitochondrial transport like all other organelles is bimodal (l; anterograde, towards the growth cone, or retrograde, towards the cell body) and can be influenced by the overall metabolic state of the neuron^31^. Within their respective motile mitochondria population, long and short axons show similar size of mitochondrial fractions moving retrograde and anterograde **(Figure 5E)** but dissecting these movements in populations across the different compartments of long and short axons, shows a trend towards increased retrograde transport in the distal compartment of long axons **(Figure 5F)**. In summary, we have found a direct correlation between mitochondrial content and axonal length in long axon using LAM arrays, indicating that mitostatic processes adapt dynamically to the increasing cell size.

**Figure 5:**
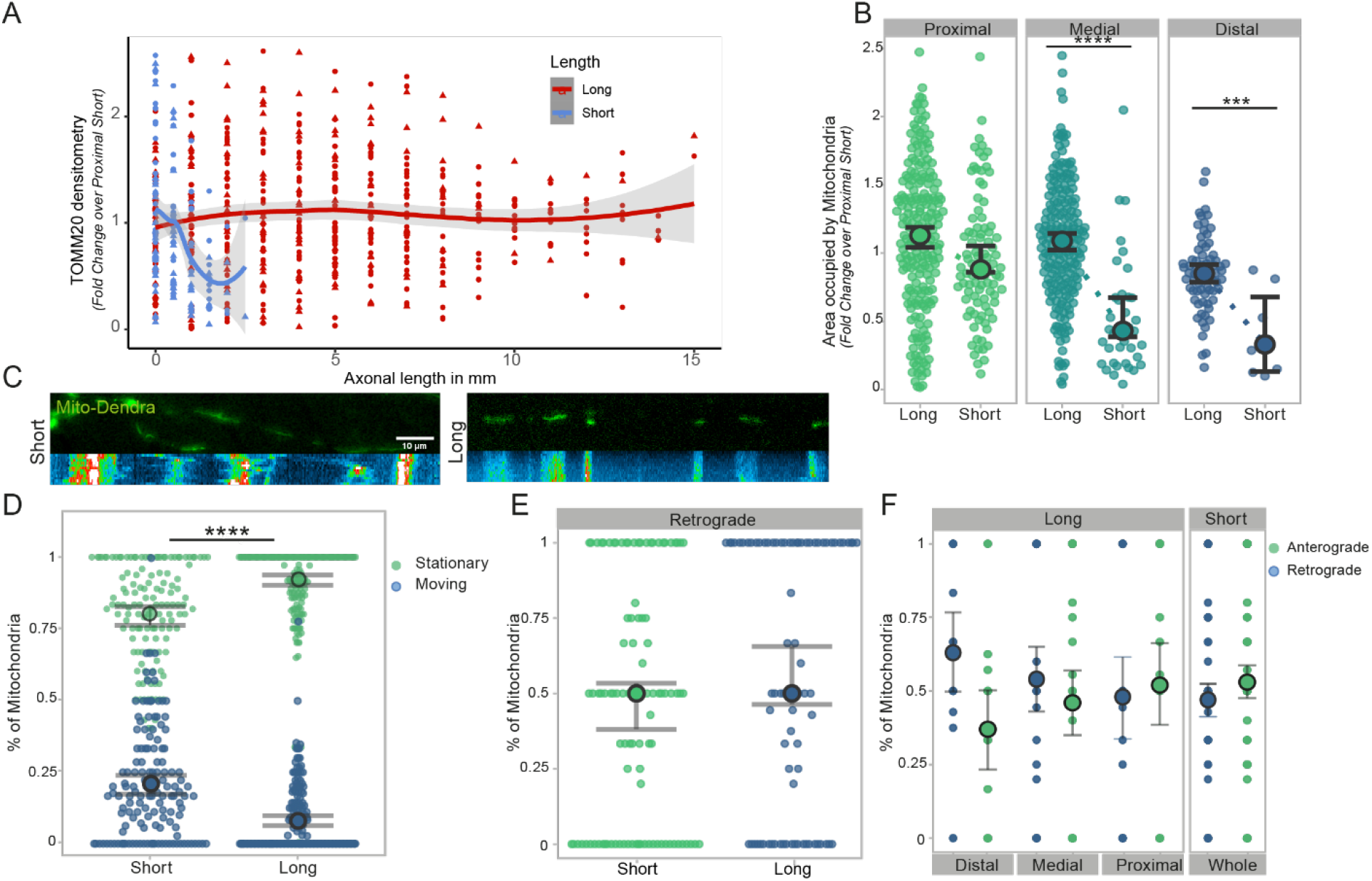
Mitochondrial content and transport are altered within long axons on LAM arrays. **(A-B)** Mitochondrial density is increased in long motor axons cultured on LAM arrays compared to short. Quantification of TOMM20 (mitochondrial outer membrane marker) using the axonal area covered by mitochondria and normalized to the proximal short compartment (A) and mitochondrial occupancy across different compartments (Mann-Whitney, ***p=0.0002, ****p<0.0001, 2 independent experimental blocks with 2 cell lines each with 3 technical replicates and 1 independent experimental block with 1 cell lines each with 3 technical replicates). **(C)** For tracking mitochondrial movement, cells were stably transfected with a Mito-Dendra2 PiggyBac system and cultured on bioengineered substrates. Fluorescent images of Mito-Dendra2 transfected motor neuron axons of different lengths and respective reconstruction of mitochondrial movements using kymographs. **(D)** Live cell imaging of Mito-Dedra2 every 10s for 2minutes was used to analyze mitochondrial shuttling along short and long axons. Quantification of mitochondrial movement shows that significantly more mitochondria are stationary compared to short motor axons (Welch’s t-test, ****p<0.0001, 1 independent experimental block with 2 cell lines each with 3 technical replicates). **(E-F)** Quantification of mitochondrial movement directionality in long and short axons cultured on LAM arrays using the fraction of moving mitochondria which are transported retrograde (Welch’s t-test, ns.) (E). Directionality of moving mitochondria, retro or anterograde (towards the growth cone), separated into compartments (F) (Welch’s t-test n.s., (E-F) 1 independent experimental block with 2 cell lines each with 3 technical replicates.

### 2.4 Axonal length affects mitochondrial morphology in a compartment specific manner

Our analysis so far has uncovered length-dependent differences in axonal mitochondrial transport and distribution, which are two of the most fundamental processes regulating axonal metabolism. However, mitochondrial shape is an important indicator of their functionality, with calcium storage capacity or reactive oxygen species generation showing clear differences between mitochondria with different morphologies^32^. Also, axonal mitochondria are in general smaller than dendritic ones, indicating a specific shape to function relationship in different cellular components^33,34^. We therefore wanted to further investigate the effect of axonal length on mitochondria by assessing potential changes in their morphology across different compartments on the LAM arrays, comparing long and short neurons. We first performed confocal imaging of both long and short neurons stained for TOMM20 and β-3 tubulin and then used a semi-supervised machine learning based approach (Ilastik) to segment TOMM20^+^ objects and classify them based on their shape descriptors into two categories: “circular” or “elongated” (**Figure 6A**). This morphological dissection of the axonal mitochondrial populations showed that while the proximal mitochondria of both long and short axons were similar, in the distal compartment long axons have significantly more circular mitochondria **(Figure 6B&C)**. Moreover, using the same dataset, we were also able to further analyze the morphology within the elongated mitochondria populations, and confirmed that even within the elongated mitochondria, the distal compartment of long axons shows a distinct morphology with less elongated branches compared to proximal mitochondria of the same group in both of long and short axons **(Figure 6D)**. Collectively these findings further support the notion that the distal segment of long axons presents specific adaptations and changes to its functional profile, with mitochondria that more resemble the axonal mitochondria described *in vivo*, when compared with the same structure in short axons or even with their own proximal counterparts, reinforcing the notion of a potential internal “threshold length” in axons that determines a rearrangement of metabolic homeostatic processes.

**Figure 6:**
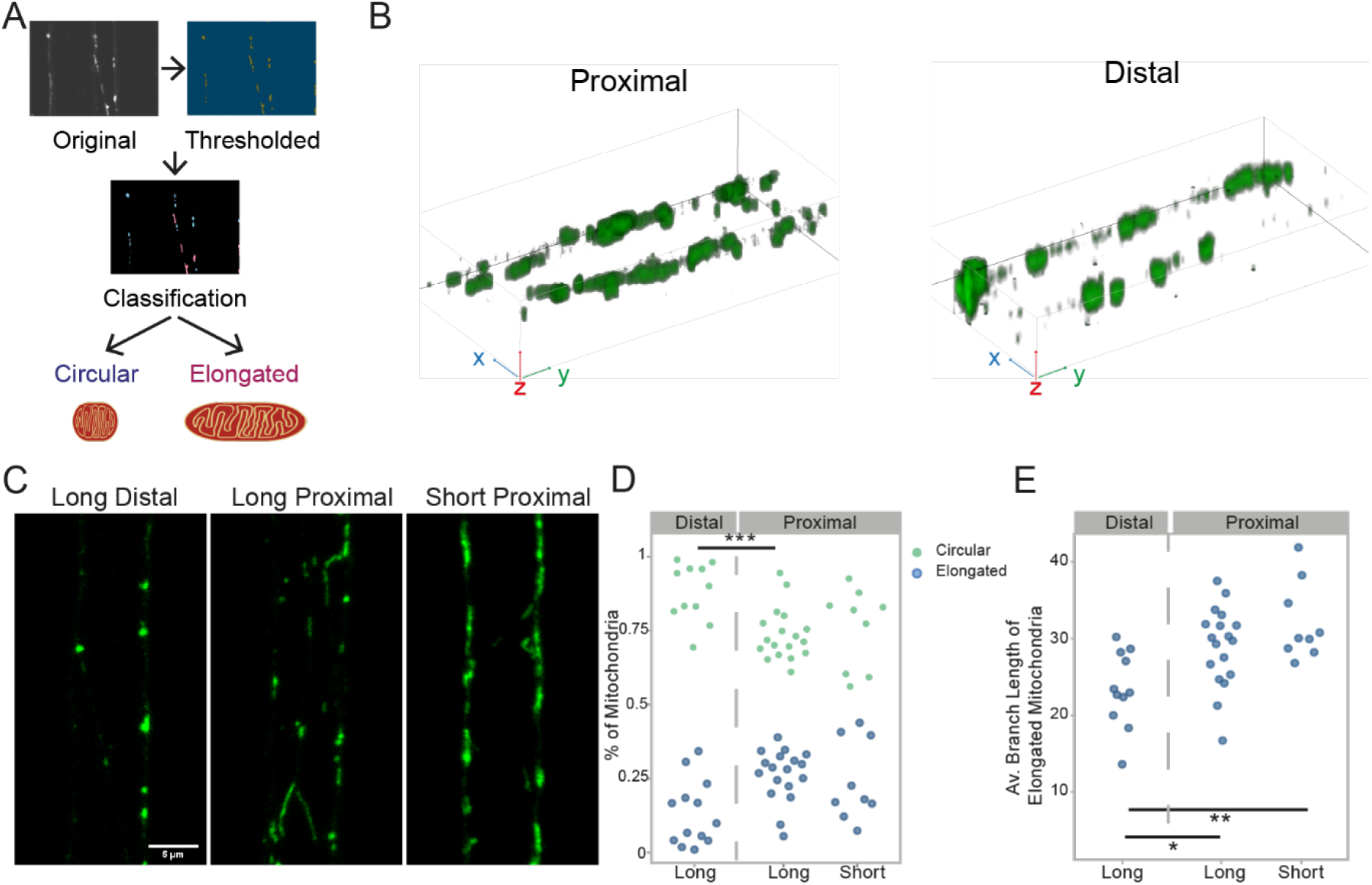
Evidence of length-dependent changes in distribution of mitochondrial morphologies within human motor axons. **(A)** Workflow of mitochondrial shape analysis using Ilastik. Object shapes were classified into “circular” or “elongated” using pixel and object classification. **(B)** 3D reconstruction of representative confocal images of proximal and distal mitochondria stained for TOMM20 in long axons on LAM arrays **(C)** Representative confocal images of TOMM20 stained mitochondria in long and short axons using LAM arrays, showing different mitochondrial morphologies in long and short axons and within long axon compartments. **(D)** Long axons show a compartment specific effect of mitochondrial morphology distribution with more circular mitochondria found in the distal compartment of long motor axons. Analysis of mitochondrial shape classification in different compartments of long and short axons (Kruskal-Wallis test with Dunn’s Multiple comparison, *** p=0.0006, 1 independent experimental block with 2 cell lines each with 3 technical replicates). **(E)** Mitochondrial object segmentation is used to calculate the average branch length of elongated mitochondria using Ilastik showing differences between long and short axons and within long axons. (One-way Annova, *p=0.0198, **p= 0.0015, 1 independent experimental block with 2 cell lines each with 3 technical replicates

### 2.5 LAM Arrays allow in-depth molecular analysis of mitochondrial dynamics and compartment specific next-generation sequencing of axonal mtDNA

Having shown that mitochondrial morphology, which is mostly shaped through fission and fusion^33^, is altered in the distal compartment of long axons, we then wanted to investigate whether these dynamics are changed in a length- or compartment-dependent way, by comparing the distribution of major fission and fusion regulator proteins in long and short axons cultured on LAM arrays. We therefore looked at Mitofusin 1 and 2 (MFNs), which are principal regulators of mitochondrial fusion and tethering of mitochondria to the ER^34 32^, as well as at Dynamin-related protein 1 (DRP1), which is recruited to the mitochondrial membrane for fission^35,36^, as marker for this analysis. After immunolabeling, we performed a densitometric analysis and used the ratio between the relative abundance of these proteins as an indication of the occurrence of fission and fusion. At the whole axon level, we found that the ratio between DRP1 and MFNs is increased significantly in long axons compared to short ones, indicating a shift towards fission **(Figure 7A)**. This finding is consistent with the observed length-dependent changes in mitochondria morphology we observed (Figure 6) and it represents further evidence on the effect of length on human axonal biology, but also -importantly-is supported by prior findings in primary mouse neurons from other groups ^35^. Moreover, upon grouping the dataset into the different compartments, this difference is most evident in the medial and distal axons **(Figure7B-C)**, in a similar pattern shown for the mitochondrial morphologies investigated above (Figure 6).

**Figure 7:**
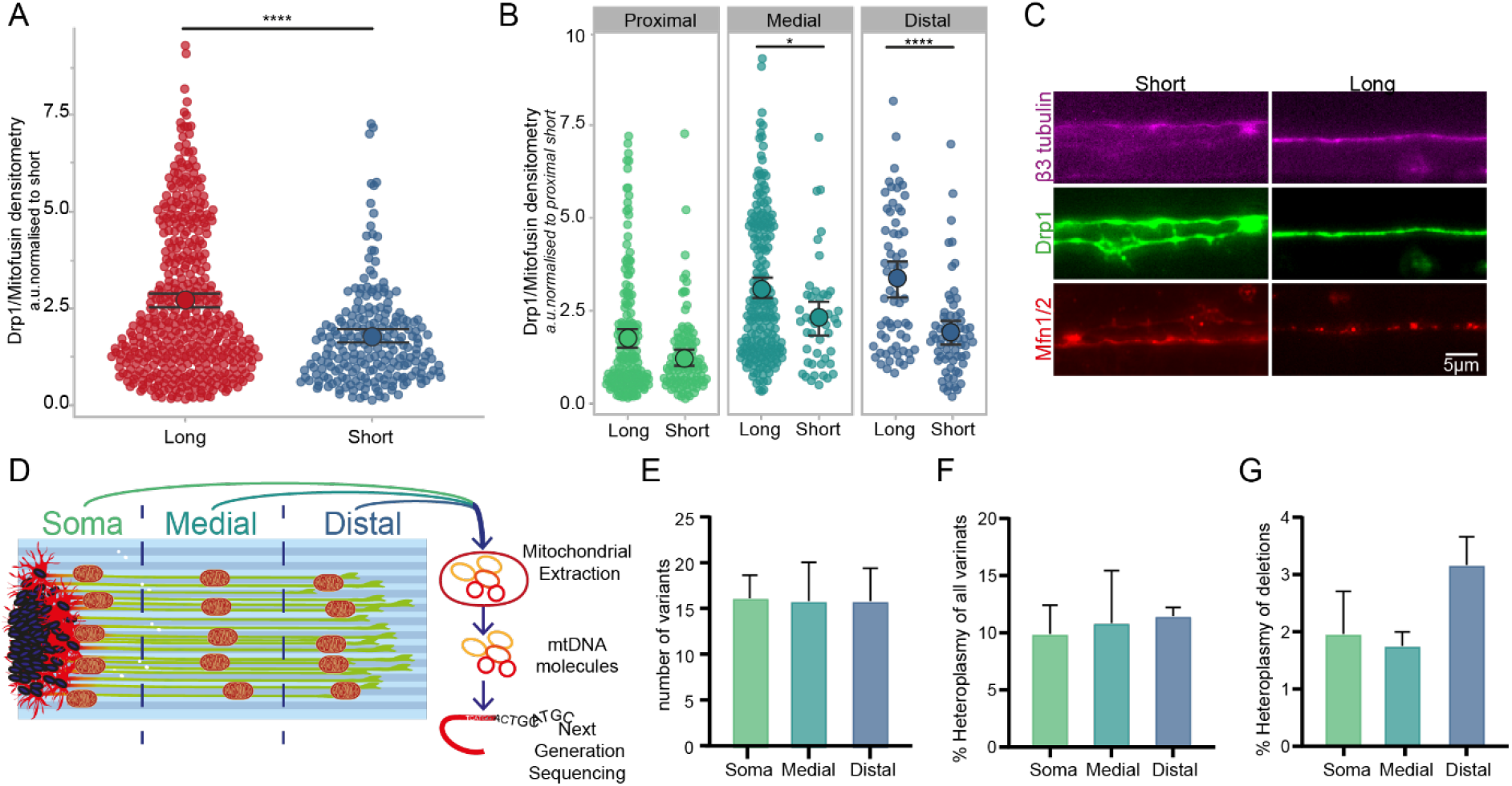
LAM array length-dependent analysis indicates alterations in mitochondrial dynamics and allows for mitochondrial genotyping within axons. **(A-C)** Long axons on LAM arrays show an altered ratio between major mitochondrial dynamics markers, DRP1 (fission) and MFNs (fusion) compared to short axons. DRP1/MFN ratio is normalized to the proximal segment of short axons (A) (Mann-Whitney, **** p<0.001). Analysis of the DRP1/MFN ratio in the compartments of long and short axons, shows that the proximal segments of short and long axons have a similar fission/fusion balance, whereas the medial and mostly the distal segment show significant alterations in the ratio leaning towards fission (B) (Mann-Whitney, *p= 0.0124, **** p<0.001). Representative fluorescent images of long and short axons for DRP1, MFN1/2 and an axonal marker, β-3 tubulin (C) 2 independent experimental blocks with 2 cell lines each with 3 technical replicates, and 1 independent experimental block with 1 cell line each with 3 technical replicates **(D)** Schematic overview showing the protocol for compartment specific next generation sequencing of mitochondrial DNA using LAM arrays. Arrays are separated into 3 compartments, soma, medial and distal, then the material in each compartment is lysed and mitochondrial DNA is extracted using silica columns. The extracted DNA is then used for next generation sequencing. **(E-G)** Mitochondrial DNA in long axons cultured on LAM arrays show compartment specific alteration with the distal compartment having an increase of large deletions in the genome. Analysis of next generation sequencing of compartment specific mitochondrial DNA looking at number of variants (E), the heteroplasmy of variants (F) and heteroplasmy of deletions >1kb. 1 independent experimental block with 1 cell line each with 3 technical replicates (Kruskal-Wallis test with Dunn’s multiple comparison).

Mitochondrial fission and fusion influence different mitochondrial morphologies, but also influence the exchange and distribution of mitochondrial DNA (mtDNA) between mitochondrial populations. Thanks to the directionality and absence of separating chambers in the LAM arrays compared to other culture systems, we were able to investigate the distribution of mtDNA by next generation sequencing (NGS) in a compartment specific manner. LAM arrays were used to culture long axons (1 cm in length), which we were then able to easily separate into soma, medial, distal compartments by physically sectioning the PDMS devices after initial live imaging to assess the position of the neurons within the arrays. mtDNA was then extracted from each compartment and analyzed by NGS to determine levels of enteroplasty by single point mutation or caused by deletions^36^ **(Figure 7D)**. The results of this analysis showed that while all compartments presented a level of heteroplasmy below the threshold of what would be considered pathological, more deletions over 1kb in size were found in the distal compartment of long axons compared to the other compartments of the same cells **(Figure 7G)**. This finding, together with the observation on DRP1/MFNs ratios reported above, would indicate that in long axons, especially in the distal compartment, the balance between fission and fusion is skewed towards relatively increased fission.

### 2.6 Axonal length determines a significant increase in axonal local protein translation

After mRNA is produced in the nucleus, peptide chains are translated at ribosomal sites, whereafter they have a limited protein-specific lifetime, during which they must be shuttled to their destination. In an elongated and tubular structure like axons, this can require significant amounts of time and local demands would not be quickly met if the translation would only happen in the soma^3,8,37,38^.

The production of local proteins within axons is key for elongation^37^, synaptic activity^38^ and other processes, but it is also directly connected to metabolism and mitochondria^39^. As we have observed numerous metabolic length-dependent phenotypes, we sought to investigate whether axonal length can also alter local protein translation.

We use a puromycin-based assay to tag newly translated peptide chains^40^, which can be then labelled with an antibody for visualization and quantification **(Figure 8A)**. Comparing long and short axons with this method reveals that long axons have significantly more active sites of local translation (identified by puromycin^+^ puncta) compared to short ones **(Figure 8B)** and that this effect is more evident moving from the proximal to the distal across compartments **(Figure 8C-E)**, indicating that length is a factor that actively influences the amount of local protein translation occurring at any given point within the axon.

**Figure 8:**
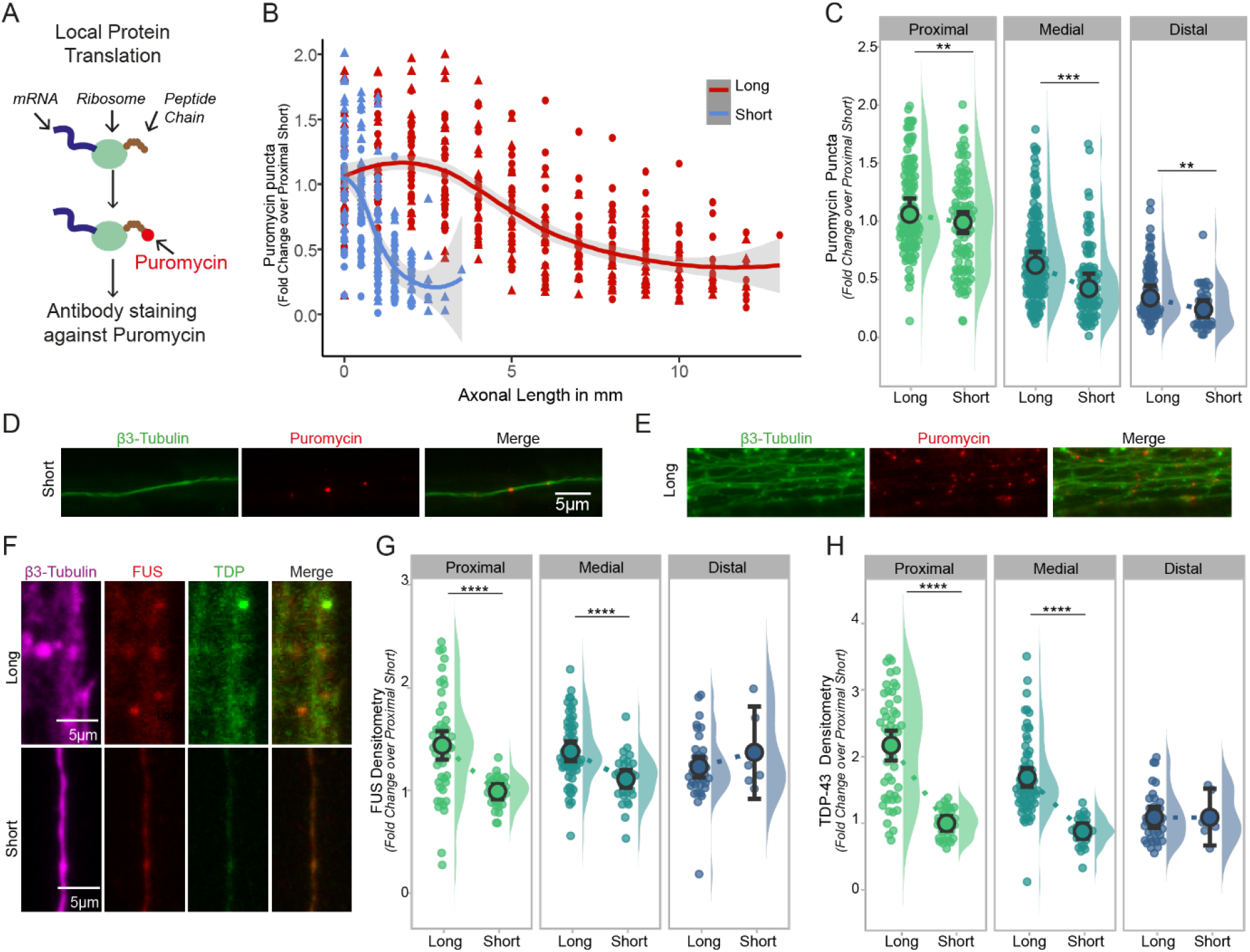
Length increases axonal local protein translation and alters distribution of key RNA binding proteins within axons. **(A)** Schematic overview of the puromycilation assay for labelling local protein translation in cells. Protein translation takes place at ribosomes which are translating mRNA into peptide chains. Puromycin is added to the living cells and labels peptide chains made at ribosomes. After fixation immunocytochemistry is used to detect puromycin which allows quantification of sides for protein translation. **(B-C)** Long axons have significantly more puromycin puncta across the whole length of the axon compared to short axons both cultured on LAM arrays. Quantification of puromycin puncta normalized over axonal total area comparing short or long MNs shows alteration in local translation at the whole axon level (B) and within specific compartments (C). (Mann-Whitney, proximal ** p= 0.0078, medial ****p<0.0001, distal **p= 0.0011) 2 independent experimental blocks with 2 cell lines each with 3 technical replicates). **(D-E)** Representative fluorescent images of puromycin labelling and an axonal marker, β-3 tubulin, showing more puromycin puncta in long axons than short axons both grown on LAM arrays. **(F)** Representative confocal images of RNA binding proteins FUS and TDP in long and short axons grown on LAM arrays. **(G-H)** Key RNA processing proteins, FUS and TDP-43, are found in the axonal compartment of long and short axons, and long axons show altered distribution of these proteins in a compartment specific manner. Quantification of the fluorescent signal of FUS (G) (Mann-Whitney, **** p<0.0001) and TDP (H) are normalized to β-3 tubulin. (Mann-Whitney, ****p<0.0001) both experiments were performed in 1 independent experimental block with 2 cell lines each with 3 technical replicates.

To maintain and regulate both translation and RNA metabolism within the axon, a number of RNA binding proteins (RBPs) are localized within the axon^41^, coordinating transport, granule formation and delivery of the mRNA to sites of local protein translation^42^, subsequently influencing the production rate of proteins at these sides. ^43^ Moreover, RBPs exhibit a broad influence on RNA splicing, stability, and transcription, wherefore it is not surprising that their malfunction makes them key player in neurodegenerative diseases^18,22,44^. We focused on two of these key RBPs, fused in sarcoma (FUS) and transactive DNA response binding protein 43 (TDP-43), as they are required in the healthy axon and are known to cause ALS^45,46^. Using LAM arrays allows us to investigate the effect of length and increased compartmentalization on location and distribution of these RBPs. Immunocytochemistry staining for FUS reveals increased intensity levels in the proximal and medial compartment **(Figure 8F-G)** and we observe a similar pattern for TDP-43, where significantly more RBP can be found in the proximal and medial compartment compared to short axons **(Figure 8H)**. This is especially important, keeping in mind that these proteins show a pathological mislocalisation^45^, for modelling neurodegenerative diseases.

## 3. Discussion

The links between cell shape and function are a fundamental area of cell biology to further explore, to better understand the molecular mechanisms that underpin homeostasis in cells that have polarized architectures like neurons.

Several techniques have emerged to probe the shape-function relationship *in vitro*, like microcontact patterning, artificial ECMs with tunable stiffnesses and several types of microfluidic devices. However, physiological relevant cell size and architecture are still very difficult to systematically model in neurons, and therefore several fundamental questions on how neuronal shape and size affects their homeostatic mechanisms and functional properties have relied on observational evidence or remained largely unexplored.

One of these questions is how neurons adapt their internal dynamics to maintain functionality across the potentially long distances that their axons can span, reaching over 1m in length in humans.

Particularly, understanding this in human neurons, has important implications for human health, as several neurodegenerative disorders may start as distal axonopathies in the periphery and travel back to the spinal cord to affect the soma in time. Critically, to shed light on these mechanisms systematically, *in vitro* modelling has several advantages, as *in vivo* experiments would be limited to developmental observations between neurons of different lengths, while the *in vitro* models not only offer the opportunity to directly study neurons but allows to directly manipulate several variables within the culture.

Our group has been interested in modelling spinal motor neurons using stem cell *in vitro* models for some time, and we developed several protocols and platforms to recapitulate different aspects of spinal cord circuitry in health and disease^10,46,47^.

One fundamental drawback of all neuronal modelling *in vitro*, and particularly of spinal motor neurons, is the fact that conventional culture systems and most microfluidic devices do not allow cells to extend axons beyond some mm in length, and in most cases confined to hundreds of μm at best, compared to cm to m scale found *in vivo*. This magnitude difference in our modelling technology, together with the ample evidence that there are several mechanisms used by neurons to regulate distal axonal processes and sense their own length, raises the question on whether the extent of axonal length *in vitro* is per se a significant factor, and whether recapitulating it *in vitro* is necessary.

To start answering these questions we first developed a bioengineered platform optimized to generate ordered arrays of human iPSC derived motor neurons, which would allow systematic investigation of length as a determinant isolated from other factors.

Our platform provides a homogenous cell environment without physical separation of compartments, which are used in conventional microfluidic systems and through this allows a dynamic generation of axonal arrays with controllable length, ranging from μm to cm in scale. While other studies have used microfluidic systems and other axonal chamber devices to force separation from soma and axons^48,49^, this is to our knowledge the first study of a simple single device system that allows to obtain human motor axons exceeding 1 cm in length from stem cell derived neurons in a reproducible and systematic way, with the possibility to clearly identify different compartments (e.g. soma, proximal and distal axons) without a multi-chamber organization that would physically separate them. At the same time, the LAM arrays allow to focus on single axons within the culture, which makes it ideal for an in-depth investigation of the effect of cell size and length on axonal homeostasis and related molecular mechanisms. Crucially, one advantage here is that -conversely to conventional axonal microfluidic devices-we do not have to vary the design of the platform to obtain different axonal length, as the axons do not grow within enclosed chambers of predefined length, but within open microgrooves. This also offers a greater degree of freedom with the analysis, as the entirety of the axon is effectively exposed to the culture medium and separation or manipulation is feasible. One example of this adaptability is the optimization of the mtDNA analysis pipeline we used in this study, where axons can be assessed by live imaging, and upon reaching the desired lengths the devices can be simply sliced in the appropriate compartments (even changing the relative dimensions of each separated compartment based on overall axonal length) and conventional extraction methods can be directly applied to process the different compartment. The same analysis in conventional microfluidic systems would have required custom designs for each axonal length, or limited the spacing between compartments, on top of also constraining the amount of material available for extraction, as only axons transmitting through open chambers could have been harvested.

Using this platform, we have been able for the first time to mechanistically characterize of the molecular and cellular effects of axonal length in human motor neurons. Importantly, our results show how cell size can regulate internal dynamics within these neurons, with longer neurons presenting significant structural and functional changes, especially within the distal portion of axons.

The first indication of these length-dependent changes was the altered NFL/NFH distribution and observed changes in calcium signaling dynamics observed in long distal axons. Cell type and functional alterations to NF networks and other axonal cytoskeletal components have been extensively studied^50^ and differences in ratio between different components of the NF network have been reported with *in vivo* studies^51,51^, but this is the first indication of length dependent alterations *in vitro* within human motor axons. Interestingly, generally NF have been associated with axonal caliber more than length, and although we have not systematically looked at axonal diameter in our system, we do not observe overt alterations comparing long and short neurons within our platform. Undoubtedly, axonal cytoskeletal organization is fundamental and extremely complex, with more details emerging constantly. For example, a recent study has indicated that axons are supported by periodic actin-spectrin ring structures^52^, which have compartment specific associations with other axonal specification factors, and play a significant role in axonal mechanical properties^53^. A more comprehensive analysis of the axonal cytoskeletal composition needs to be performed comparing long and short human axons, potentially using super-resolution microscopy to uncover differences in ultrastructural organization. The alterations observed in cytoskeletal components were matched by alterations in the frequency and amplitude of calcium waves propagating because of spontaneous activity. Although there is scope to further validate our functional calcium imaging analysis with more sensitive electrophysiological characterization, interestingly both lines of evidence are consistent within our data in indicating the same approximate distance from the soma for length-dependent changes. Moreover, several of the observed length-dependent alterations are mechanistically linked: for example, Ca^2+^ metabolism is tightly linked to mitochondrial dynamics and function, and both spike frequency and mitochondrial dynamics and morphology are consistently altered in distal portions of long axons. Also, calcium waves and other wave-based signaling have been proposed as a potential sensing mechanisms for neurons to gauge their own size past the mm scale^2^, which would fit with our observations.

Strikingly, both mitochondrial populations and local protein translation are altered by axonal length in a homogeneous way along the axon, conversely to NFs, calcium and ER distribution which change progressively, potentially indicating separate “long” and “short” phenotypes that are enacted upon reaching a certain size. The finding that ER shape and contact sites are linked to hereditary spastic paraplegia, a disease which is mostly prominent in long motor axons, also indicates specific adaptations of molecular mechanisms to the axonal length. These observations would be consistent with the model of a longer axons having a consistently higher demand for ATP production, transport and local production of proteins, and therefore having to adapt by increased homeostatic functions as ATP has a limited diffusion capacity.

Using our LAM arrays we found a skewed balance between fission and fusion, which has been indicated in a previous study^35^, and we also observed a trend towards increased deletions in the distal mtDNA and the increased retrograde transport. We could speculate that one possible scenario for these observation could be that in the terminals of long axons there could be a slower mitochondrial turn-over, and that alterations in fission and fusion dynamics help with the maintenance of a healthy damaged DNA can be removed, although more focused investigation is certainly needed for a definitive answer.

When systematically evaluating axonal length within our platform, one potential confounding variable could be the fact that “long” and “short” neurons are grown for different amount of time within the LAM arrays before analysis, and that could translate to maturation differences which would supersede length-dependent effects. For this reason, we have taken in this study all possible attention to this factor and performed additional experiments where we match time in culture across our platform, PDMS without microtopography and conventional tissue culture plates (Figure S4), and we show that i) LAM arrays still allow to obtain the longest neurons in comparison and ii) even accounting for maturation differences the mitochondria phenotypes seem to be exclusively linked to axonal length. Moreover, there is some variability in the time necessary for MNs on LAM arrays to extend axons to 1cm (between 2 and 4 weeks) but even accounting for this, at the analysis stage length-matched cultures of different age display a consistent phenotype.

Ultimately, this study presents indications for axonal length directly influencing internal homeostatic mechanisms within neurons and suggest a length-dependent switch of homeostasis and distinct phenotypes, which we have defined as “short” and “long”. Of course, this definition of long and short is based on our initial characterization and it is in no way an exhaustive or fully comprehensive analysis, which future studies will conduct varying axonal length and parsing the axon mm by mm.

However, the hypothesis of a “threshold length” that determines a switch between different mechanisms of maintenance traces back also to the idea that maintaining homeostasis within a long axon can create “bottlenecking” of resources when demands at the terminal exceed the transport capacity, and therefore below certain lengths a lot of adaptations evolved to maintain this delocalized biology are in fact not necessary. For example, Le Masson and coworkers ^54^ have shown computationally in the context of ALS that while Ca2+ dependent hyperexcitability phenotypes in MNs can be created in the axon terminals by imbalances of ATP that are not promptly resolved, they do not actually generate a propagating wave of hyperexcitability that reaches the soma unless the axon extend past a certain length, in short “bottlenecking” the potential diffusion of resources from the soma.

As a detailed mechanism for length sensing in neurons that works past the mm scale has not been completely established and that several MN centric diseases start as distal axonopathies (with involvement of the very pathways and proteins we see changing between long and short neurons) platforms like LAM arrays, where cell shape is more faithfully recapitulated thanks to the bioengineering, offer a crucial advantage that has clear impacts for basic neurobiology and neurodegeneration research.

## 4. Conclusion

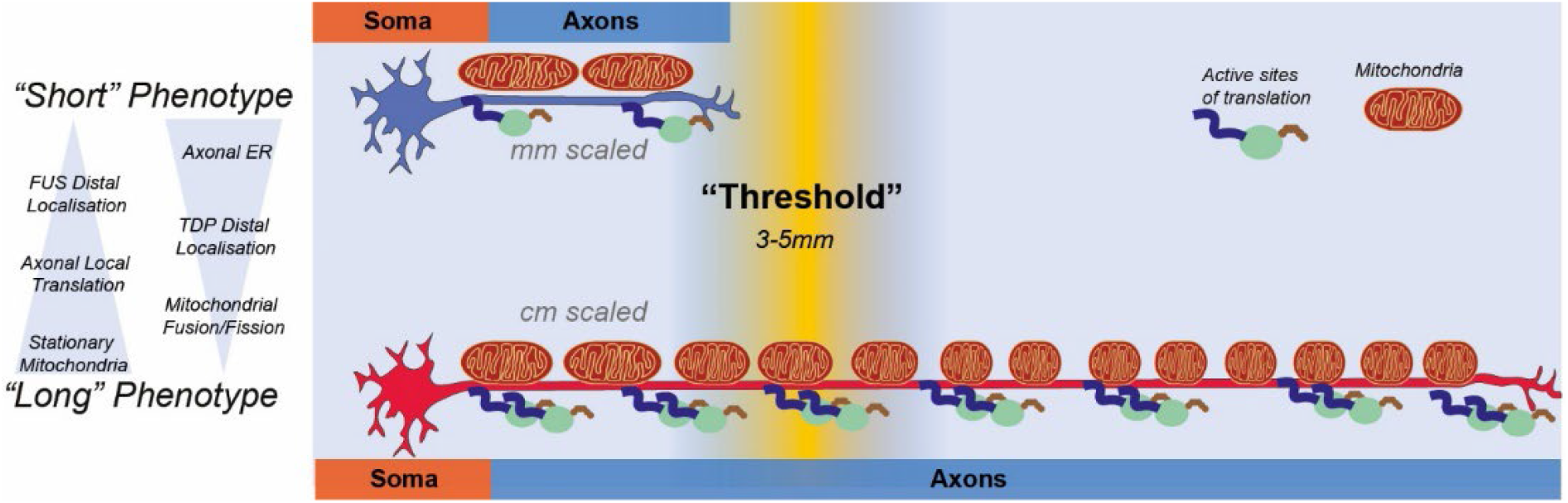

Our study breaks new ground into fundamental neurobiology by applying bioengineering principles to the study of an essential question and combining different techniques into a novel and easily reproducible platform that is ideally designed for pushing the boundary of what is achievable in the field of axonal biology.

## 5. Methods

### Constructs

The PiggyBac Mito-Dendra2 construct was designed from the Addgene plasmid #55796 (http://n2t.net/addgene:55796; RRID: Addgene_55796). Mito-Dendra2 was a gift from David Chan.The Mito-Dendra2 sequence was inserted to a PiggyBac backbone construct along with a T2A Neomycin selection cassette using the GeneArt assembly kit (Invitrogen). We also used a pgK-Puro-CMV-GFP and *pPb-CAG-RFP-Hygro* construct cloned in the lab and a PiggyBac vector containing the transposase.

### Cell culture

Cells were kept with 5% CO_2_ in humidified atmosphere at 37°C. IPSC derived control progenitors were obtained as previously described ^10^and more information can be found in Table1. These progenitors were cultured in growth medium, containing base medium 50% advanced DMEM (Gibco) and 50% NeuroBasal (Gibco) with NB27 (Gibco) and B2 (Gibco) supplements and 100ug/mL Pen-Strep (Gibco), 2mM L-alanyl-L-glutamine dipeptide (Gibco) and in growth medium additionally FGF (20ug/mL) (Gibco). MN progenitors were differentiated using base medium and additional compound E (Enzo) (0.1uM) and the growth factors BDNF (10ug/ml) and GDNF (10ug/ml). All cells were grown on Matrigel (Corning) coated plates unless stated differently.

**Table 1.**
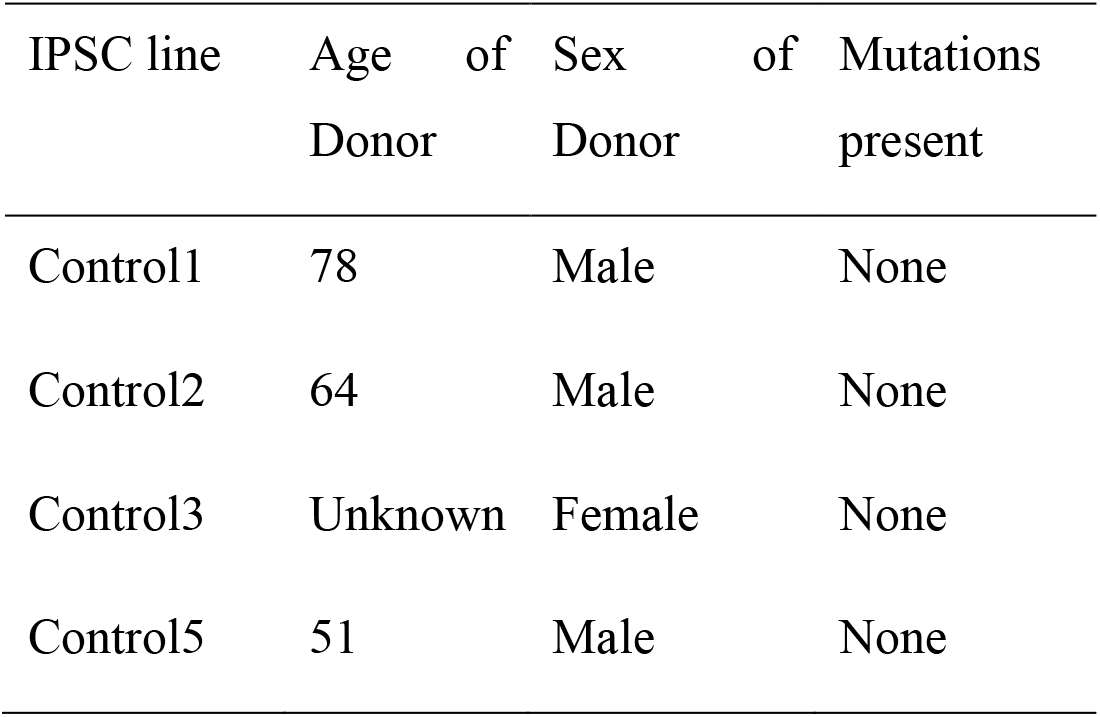
Details of iPSC lines.

| IPSC line | Age of Donor | Sex of Donor | Mutations present |
| --- | --- | --- | --- |
| Control1 | 78 | Male | None |
| Control2 | 64 | Male | None |
| Control3 | Unknown | Female | None |
| Control5 | 51 | Male | None |

For generation of MN aggregates, cells were cultured for additional 2 days after reaching confluency in growth medium. Then cells were detached using EDTA-PBS (0.5mM) (Gibco) for 3min at room temperature and pelleted. The cell pellet was dissolved in medium and transferred to a petri dish (90mm) with 10mL growth medium. Cells are then kept in suspension on an orbital shaker (Stuart) at 60 rpm in an incubator as above described. 2 days after this expansion phase, the medium is changed to basis medium for another 2days and from then on fed 2x per week with base medium. After this pre-differentiation phase, neural aggregates were either placed immediately on a cell culture substrate or up to 2 weeks later.

### Manufacturing of the model platform

Silicon masters were prepared as described before^55^ by using SU-8 2002 at 1000 rpm for 40 s on a silicon wafer on a spin coater (Polos) and then prebaked at 95°C for 2minutes. On top of the prebaked surface a 10µm wide striped pattern interrupted with plateaus every 5mm with a length of 250um photomask was aligned (Kiss MA6 mask aligner) and UV exposed. To develop the pattern the master was baked at 95°C for 300s and washed in Su8-developing liquid. For flat surfaces only the silicon wafer without SU8 was used.

For generation of micropatterned surfaces a silicon elastomer (poly dimethyl siloxane) was used (Sylgard 184, Dow Corning). The base substrate was mixed with different weight ratios of the curing agent (1:5, 1:10, 1:25) and desiccated. The mixture was spun on a spin coater using a silicon master (flat or micropatterned) at 300rpm. The covered master is cured on a heat plate for 5-10min at 100°C. Low ratios as 1:25 were kept overnight at 60 °C.

For biocompatibility, the cured PDMS was treated with oxygen plasma (30s, 100%, 7sccm) (Henniker Plasma) and coated with poly-D-Lysine 0.01% (PDL) (Gibco) for 15min and laminin (Sigma) overnight unless stated differently in the figure legends.

Neural aggregates were transferred onto the biofunctionalized substrate using a pipette. During these processes’ aggregates were accumulated in the tip and then gently placed on the substrate to prevent spreading of aggregates across the material. Subsequent culture of aggregates on the substrate was performed as described in Experimental Section-Cell Culture. For imaging of these cultures, PDMS was removed from the tissue culture plastics and placed on a glass slide and sandwiched between this and a cover slip with FluorSave mounting medium (Millipore).

## Transfection methods

### Electroporation

First, we generated stable motor neuron progenitor lines using the PiggyBac system with the mitochondrial tracking plasmid (Mito-Dendra2). Motor neuron progenitors were detached using accutase (Gibco) to generate a single cell suspension and pelleted at 800rpm for 3min. After removing the excessive accutase, the pellet was resuspended in PBS to count cells. For transfections with the plasmids, we used the Neon Pipette system (Invitrogen), with the respective neon transfection kit (Invitrogen) and manufacturer’s instructions were followed. In short, no more than 10k/μL cells were resuspended in 110uL Electrolyte buffer and both plasmids (Mito-Dendra2 and the transposase containing construct) were used at 1ug/1mio cells and added to the buffer containing the cells. The cuvette which will conduct the electroporation pulses into the pipette was filled with 3mL buffer. We then used 3 pulses of 10ms with 1000mV to electroporate our samples. Afterwards, the electroporated cells in the neon pipette were seeded into a prepared Matrigel coated plate with growth medium.

### Transfection with reagents

For analysis of axonal length in conventional culture or directionality, we generated stably expressing GFP or RFP motor neuron progenitors. Motor neuron progenitors were sparsely cultured on a 24-well plate and one day after passaging transfected using Mirus LT1(Mirus Bio) transfection reagent. For this we used per well 0.5ug of each plasmid (GFP/RFP+ transposase containing plasmid) in 200uL of pen-strep free growth medium. This solution was well mixed and added to the wells which also contained 200uL of pen strep free medium. The medium containing the transfection reagent was exchanged with growth medium after 24h. Cells were cultured to confluency and then pooled into a 6-well plate and there further expanded.

### Mitochondrial tracking

We cultured the sparsely but stably expressing Mito-Dendra2 motor neuron progenitors (Transfection Method-Electroporation) until they reached confluency and followed the above-described protocol to generate neural aggregates. Aggregates were seeded on micropatterned surface in 8well -polystyrene slides (Ibidi) or glass bottom 6-well plates (Cellvis) and grown until desired length was reached (short ∼ 5d, long ∼3weeks) Cells were imaged in NB basal medium without phenol red (Gibco) with Anti-oxidative supplement (Sigma-Aldrich) to reduce the oxidative stress. Imaging was performed using Ti2 Eclipse system with a 40x ELWD objective with 200ms exposure and 100% LED power and images were taken every 10s for 2 minutes.

### Calcium tracking

Long axons were cultured on LAM arrays and then loaded with 1uM Fluo4AM (Invitrogen) for 30min., in base medium and 100nM SiR-tubulin (Spirochrome) for 30 min then 2x wash with PBS, then base medium with compound E was added. Samples were imaged with a 20x ELWD every 100ms for 100 frames and used the SiR-tubulin to find and focus on the axons.

### Directionality analysis

Motor neuron progenitors were transfected using the transfection reagent method described above using the PiggyBac RFP plasmid. After this, cells were selected using hygromycin (500 µg/ml) (Sigma) for 48h ours to generate pure RFP^+^ cultures, which were then cultured until confluent, and medium was changed to base medium for 1day prior seeding on the different substrates. Microgrooves and flat PDMS were prepared as described above and coated with PDL and laminin. Then cells were seeded on the substrate in respective cell numbers ranging from 5000-150.000 in a 96-well plate. We continued the cell differentiation with base medium and compound E for 7 days. These cells were fixed and stained as described in the immunocytochemistry section for β-3 tubulin and DAPI and images were acquired using a fluorescent microscope. Images with microgrooves were rotated so the directionality measured is standardized in ImageJ and the β-3 signal was used to measure alignment of axons to the given topography, with the in-build line tool and plot profile in ImageJ.

### Puromycin assay

Puromycin (Gibco) was added to the medium in a final concentration of 10ug/mL together with a dissociation blocker CHX (10μg/mL) for 15min, then washed out with3x PBS. Cells were then fixed and stained with an anti-puromycin antibody for visualization. ^40^ Cells were fixed and stained for β3 tubulin as an axonal marker and DAPI for nuclear localization, and an anti-puromycin antibody to detect sides of local protein translation. These staining were imaged using widefield fluorescent microscope at 20x SWD every 1mm in long arrays and every 500um in short arrays.

### Mitochondrial shape

Samples were stained with TOMM20 and imaged on an inverted Zeiss confocal using an 63x Oil objective. Z-stacks were taken across the microgrooves and projected using the SUM function in ImageJ. These images were then used to identify mitochondria in a machine learning based program, Ilastik.^56^ Here we used a combinatorial approach of pixel and object classification in one pipeline. 3D reconstructions were generated in ImageJ using the volume viewer plugin.

### Immunostaining

Cells were fixed using 4% PFA (Boster) for 15 min. Afterwards they were washed 3x with PBS. Cells were permeabilized with 0.1% Triton (Sigma) in PBS for 10min at room temperature. After washing the blocking solution containing 3% goat serum (GS) (Sigma) was added for 30min at room temperature. Antibodies (AB) were diluted in 0.05% Triton 1.5% GS in PBS. Antibodies and their concentration can be found in **Table2**. All primary AB were incubated for 1h a room temperature, then washed 3x with PBS. Following secondary AB were used: Alexa 555 goat anti-mouse, Alexa 488 goat anti-rabbit, Alexa 647 goat anti-mouse, Alexa 488 goat anti-mouse, Alexa 405 goat anti-mouse (all Invitrogen). They were diluted in the same solution as primary AB. All secondary AB were used with a final concentration of 2ug/mL. The secondary AB was incubated for 30min at room temperature and then cells were washed 3x with PBS. Only FUS and TDP-43 staining were performed using 0.5% Triton in PBS and instead of goat serum we used bovine serum albumin and the primary antibodies were incubated overnight. Images were taken for DRP1/MFN experiments at 63x Oil immersion on a fluorescent microscope, TOMM20, calreticulin and NFL/NFH densitometries were calculated based on 20x short working distance images taken at a fluorescent microscope.

**Table 2.**
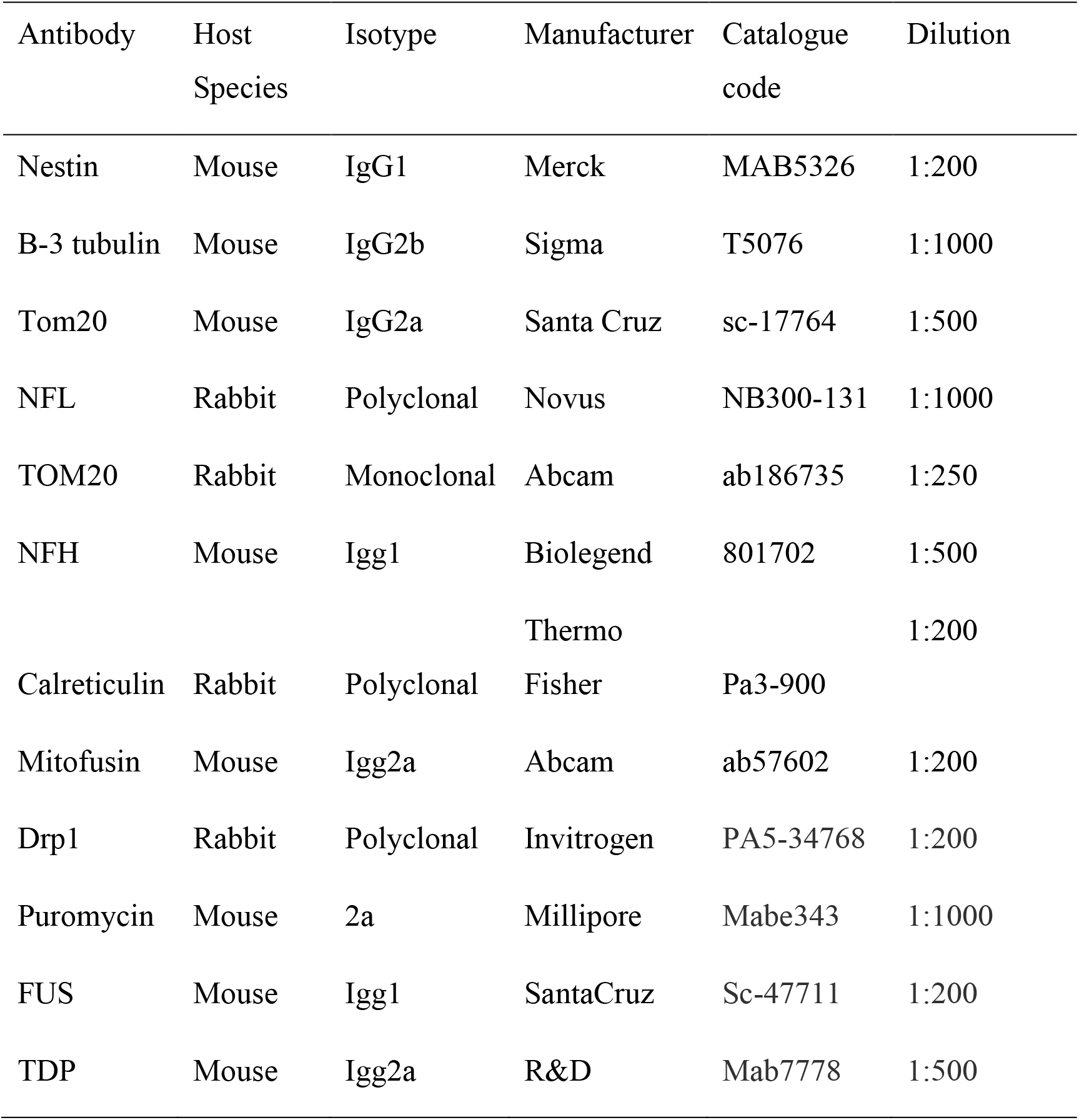
Antibodies

### Image Analysis

For mitochondrial dynamics as a first step the fluorescence of each stack was normalized using the NeuroCyto plugin (Normalize Movie). Time laps of Mito-Dendra2 positive axons were processed with the KymoResliceWide ImageJ tool. A line with a width of 10um was used to track the mitochondria. For generation of a Kymograph the maximum intensity projection was used. This Kymograph was analyzed using the NeuroCyto plugin (Analyze Kymograph). For immunocytochemistry experiments, Ilastik was used to identify the axonal object as a mask and nuclei and in some cases debris. These masks were used in Cellprofiler ^57^ to measure intensities of the staining only in axons. The resulting values were then normalized to the background signal and the area. For measurements signal was normalized to β 3 tubulin unless a ratio between two proteins of interest were calculated.

### Statistical analysis

Generally, the data was tested for normality (Shapiro-Wilk) and then treated accordingly. For nonparametric tests with 2 samples the Mann-Whitney test was used. For comparison of more than 2 parametric samples an ordinary one-way ANNOVA was performed with Turkey’s multiple comparisons. For nonparametric analysis a Kruskal-Wallis test with Dunn’s multiple comparison was used. Graphs were designed using custom R files or the SuperPlotsOfData app.

### Long range PCR for mtDNA analysis

The forward and reverse primers used for one amplicon PCR were the following: 16426Fw (5′-CCGCACAAGAGTGCTACTCTCCTC-3′) and 16080Rv (5′-TGTTGATGGGTGAGTCAATA-3′. PCR was performed using the LongRange PCR Enzyme Mix (QIAGEN®) and 5 ng of DNA isolated from cultured long motor neurons on LAM arrays, an initial 3-min incubation at 93 °C was followed by 30 cycles of PCR with 30 s of denaturation at 93 °C, 1 min at 58 °C of annealing and 17 min of extension at 68 °C; the reaction is completed by 1 cycle of final extension at 68 °C for 20 min. 5 μL of PCR product was analyzed on 1% agarose gel with GeneRuler 1kb DNA ladder (ThermoFisher).

### Library preparation for Illumina sequencing of mtDNA

PCR products were diluted to a final concentration of 0.2 ng/μL and processed according to the Nextera XT DNA Library Prep protocol (Illumina). Indexed DNA libraries were pooled together with equal molar ratios and sequenced on MiSeq platform with a v3 Illumina Flowcell (600 cycles) and a paired-end read chemistry.

### mtDNA sequencing analysis

Reads generated were analyzed through two pipelines

1. For heteroplasmy analysis, reads generated were mapped to the revised Cambridge Reference Sequence (rCRS) using the dedicated application on BaseSpace website (Illumina®) called mtDNA Variant Processor; then aligned reads were analyzed for variants calling and heteroplasmy levels detection with mtDNA Variant Analyzer tool. Analysis parameters were set as follows: Minimum Basecall Quality Score at 30, Analysis threshold at 1%, Interpretation threshold at 1%, Minimum read count at 10x. The output generated by the Analyzer tool is used as input data for the HaploGrep tool (https://haplogrep.i-med.ac.at) for the calculation of the mitochondrial haplogroup. The average percentage of single point heteroplasmy was then calculated for each sample, excluding homoplasmic variants associated with the haplogroup.
2. For mtDNA deletions analysis, reads were mapped to the rCRS reference genome using the BBmap tool (sourceforge.net/projects/bbmap, version 38.86), a dedicated software for the alignment of genomes with large indels. In order to calculate the percentage of heteroplasmy of mtDNAs affected by the alteration, the number of reads containing the deletion were normalized over the number of normal reads aligned, for each mitochondrial genome position, then the average percentage of heteroplasmy was calculated.

## Acknowledgements

The authors wish to acknowledge the support of the UK Biotechnology and Biological Sciences Research Council [BB/T011572/1], the Royal Society [RSG\R1\180114] and the Wellcome Trust [213949/Z/18/Z]. This research was funded in whole, or in part, by the Wellcome Trust. For the purpose of Open Access, the author has applied a CC BY public copyright licence to any Author Accepted Manuscript version arising from this submission.

## Supporting Information

**Figure S1:**
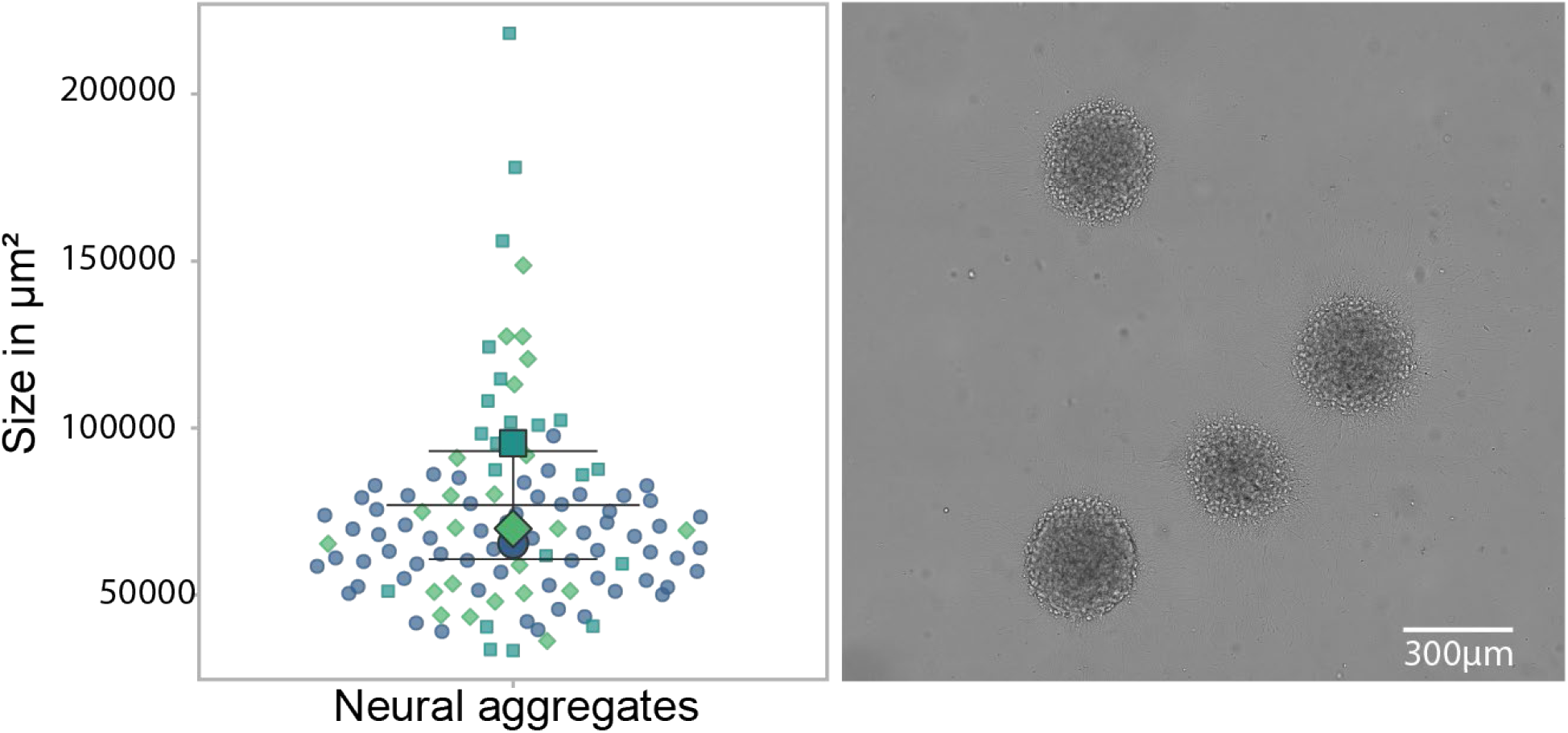
Size of neural aggregates is homogenous across experiments. We cultured neural aggregates as (see Experimental section. Cell culture) and placed them on culture substrates. After 1 day in culture, we measured the area occupied by a single aggregate. 1 experimental block with 3 hIPSC derived cell lines and 3 technical replicates.

**Figure S2:**
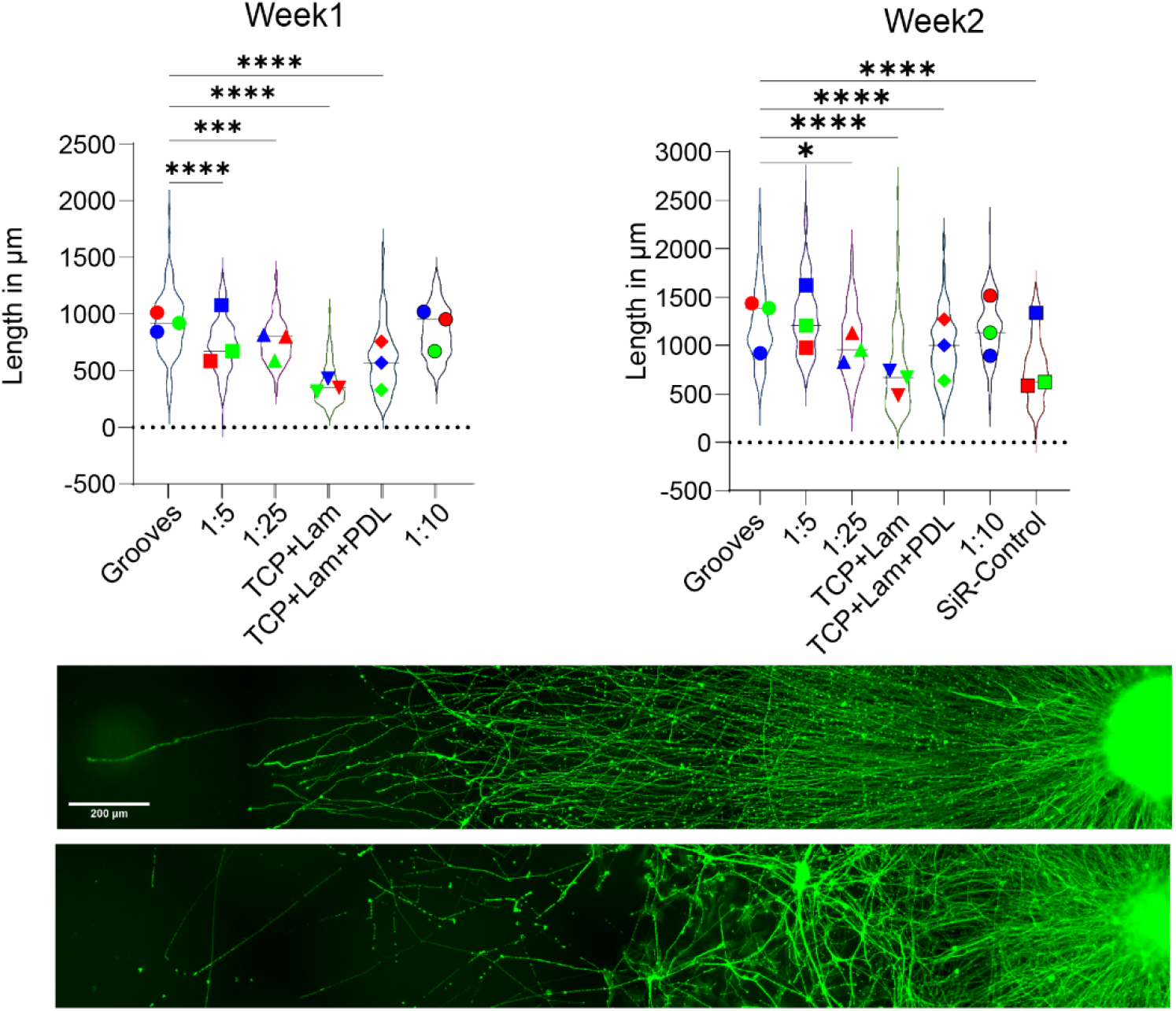
Influence of substrate properties on axonal elongation. We tested different PDMS stiffnesses and coatings to optimize axonal outgrowth. The best results were produced across time with 1:10 PDMS and microgrooves, using 1:10 PDMS with soft-litography Representative images show the effect of suitable substrate on axonal elongation, with more homogenous growth. 3 experimental block with 1 hIPSC derived cell lines and 2 technical replicates.

**Figure S3:**
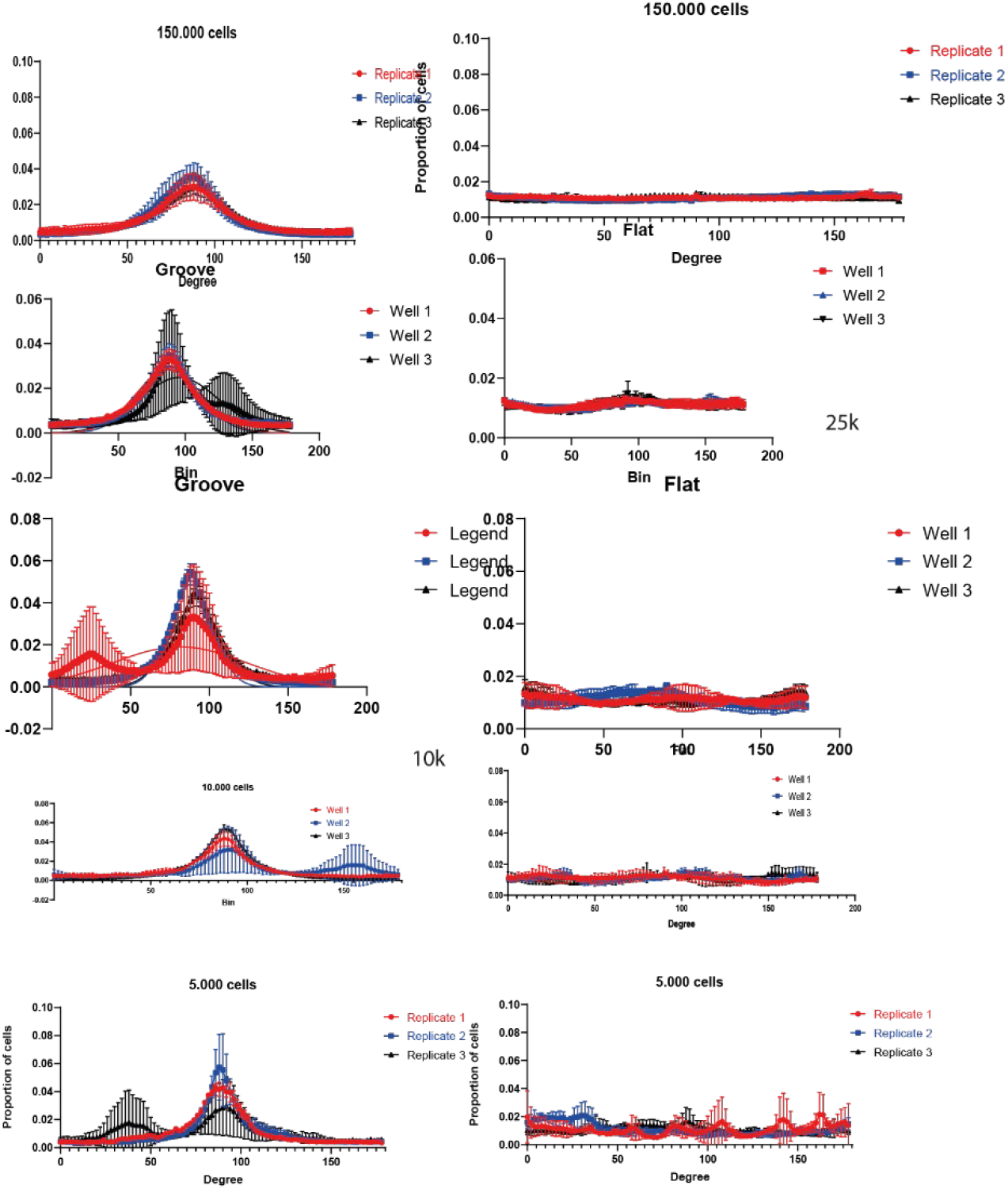
Microtopography is sufficient to align cells across different densities. Microtopography is sufficient to align cells up to 150.000 cells in a96-well plate but with lower densities chemotaxis is counteracting this effect an cells are less aligned. On flat substrate no alignment can be found. For these experiments we used RFP motor neurons progenitors cultured for one day in base medium and then continued differentiating on the respective substrates.

**Figure S4:**
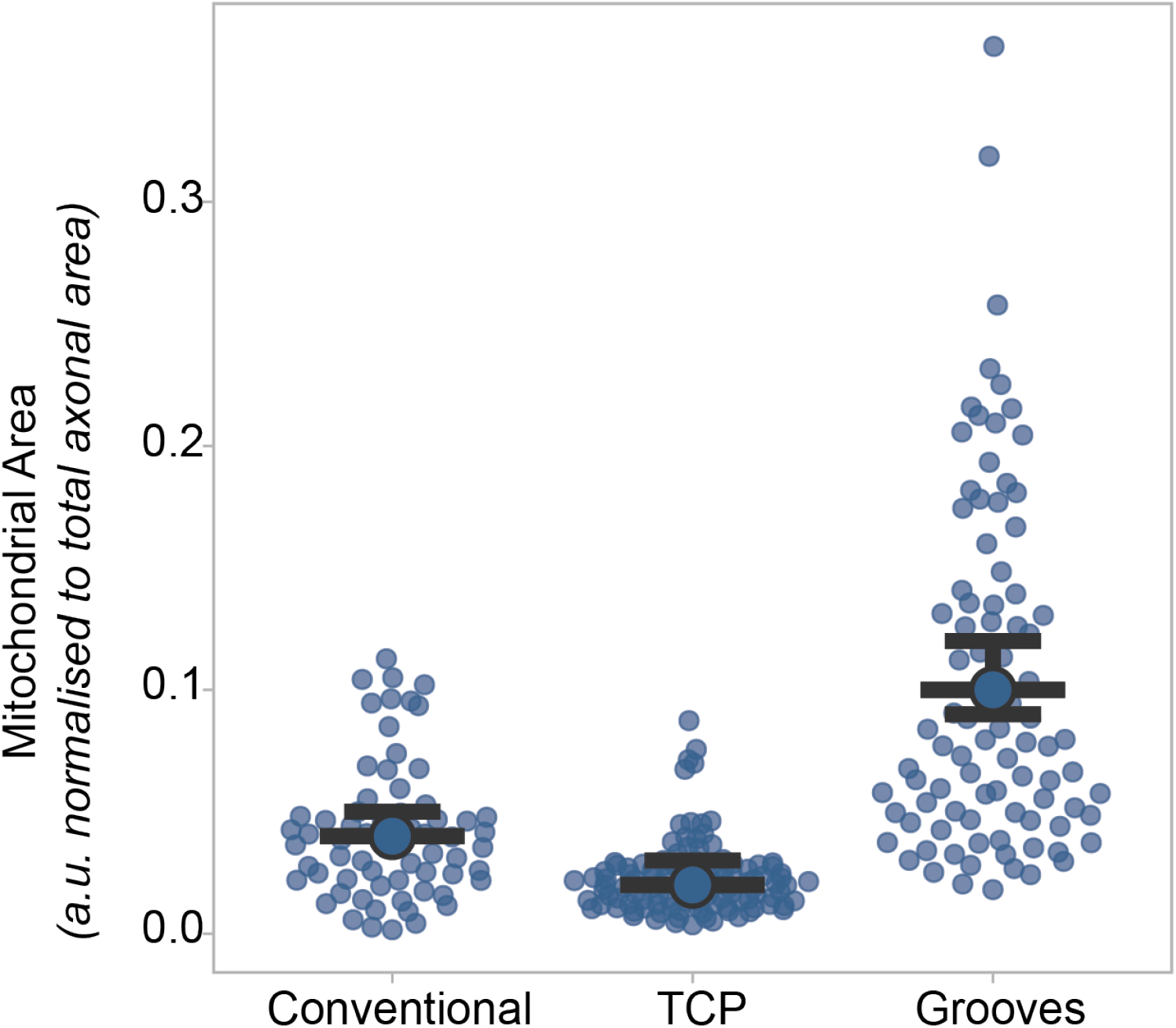
Dissection of axonal length from age of culture. Cells were cultured on microgrooves or tissue culture plastics as neural aggregates, and as single cells partially GFP+ on tissue culture plastics, called conventional culture to observe axonal outgrowth and measure mitochondrial content. Even tough, time in culture was the for all cultures 3 weeks, we observe changes in mitochondrial content, with increased populations on microgrooves. In Figure 2 the respective axonal length can be found, indicating a length dependent effect.

## References

1 Misgeld, T. & Schwarz, T. L. Mitostasis in Neurons: Maintaining Mitochondria in an Extended Cellular Architecture. Neuron 96, 651–666, doi:10.1016/j.neuron.2017.09.055 (2017).

2 Albus, C. A., Rishal, I. & Fainzilber, M. Cell length sensing for neuronal growth control. Trends Cell Biol 23, 305–310, doi:10.1016/j.tcb.2013.02.001 (2013).

3 Terenzio, M., Schiavo, G. & Fainzilber, M. Compartmentalized Signaling in Neurons: From Cell Biology to Neuroscience. Neuron 96, 667–679, doi:10.1016/j.neuron.2017.10.015 (2017).

4 Sabry, J., O’Connor, T. P. & Kirschner, M. W. Axonal transport of tubulin in Ti1 pioneer neurons in situ. Neuron 14, 1247–1256, doi:10.1016/0896-6273(95)90271-6 (1995).

5 Chamberlain, K. A. & Sheng, Z. H. Mechanisms for the maintenance and regulation of axonal energy supply. Journal of Neuroscience Research 97, 897–913, doi:10.1002/jnr.24411 (2019).

6 Lin, M.-Y. et al. Releasing Syntaphilin Removes Stressed Mitochondria from Axons Independent of Mitophagy under Pathophysiological Conditions. Neuron 94, 595-610.e596, doi:10.1016/j.neuron.2017.04.004 (2017).

7 Terenzio, M. et al. Locally translated mTOR controls axonal local translation in nerve injury. Science 359, 1416–1421, doi:10.1126/science.aan1053 (2018).

8 Costa, I. D. et al. The functional organization of axonal mRNA transport and translation. Nat Rev Neurosci 22, 77–91, doi:10.1038/s41583-020-00407-7 PMID - 33288912 (2021).

9 Galiano Mauricio R. et al. A Distal Axonal Cytoskeleton Forms an Intra-Axonal Boundary that Controls Axon Initial Segment Assembly. Cell 149, 1125–1139, doi:10.1016/j.cell.2012.03.039 PMID - 22632975 (2012).

10 Hall, C. E. et al. Progressive Motor Neuron Pathology and the Role of Astrocytes in a Human Stem Cell Model of VCP-Related ALS. Cell Rep 19, 1739–1749, doi:papers3://publication/doi/10.1016/j.celrep.2017.05.024 (2017).

11 Magrané, J., Cortez, C., Gan, W.-B. & Manfredi, G. Abnormal mitochondrial transport and morphology are common pathological denominators in SOD1 and TDP43 ALS mouse models. Hum Mol Genet 23, 1413–1424, doi:papers3://publication/doi/10.1093/hmg/ddt528 (2014).

12 Palau, F., Estela, A., Pla-Martin, D. & Sanchez-Piris, M. The role of mitochondrial network dynamics in the pathogenesis of Charcot-Marie-Tooth disease. Adv Exp Med Biol 652, 129–137, doi:10.1007/978-90-481-2813-6_9 (2009).

13 Freeman, O. J. et al. Metabolic dysfunction is restricted to the sciatic nerve in experimental diabetic neuropathy. Diabetes, db150835, doi:10.2337/db15-0835 (2015).

14 Mattedi, F. & Vagnoni, A. Temporal Control of Axonal Transport: The Extreme Case of Organismal Ageing. Front Cell Neurosci 13, 393, doi:10.3389/fncel.2019.00393 (2019).

15 Maimon, R. et al. A CRMP4-dependent retrograde axon-to-soma death signal in amyotrophic lateral sclerosis. EMBO J, doi:10.15252/embj.2020107586 (2021).

16 Rotem, N. et al. ALS Along the Axons – Expression of Coding and Noncoding RNA Differs in Axons of ALS models. Sci. Rep. 7, 44500, doi:papers3://publication/doi/10.1038/srep44500 (2017).

17 Stoklund Dittlau, K. et al. Human motor units in microfluidic devices are impaired by FUS mutations and improved by HDAC6 inhibition. Stem Cell Reports, doi:10.1016/j.stemcr.2021.03.029 (2021).

18 Harley, J., Hagemann, C., Serio, A. & Patani, R. FUS is lost from nuclei and gained in neurites of motor neurons in a human stem cell model of VCP-related ALS. Brain 143, e103, doi:10.1093/brain/awaa339 (2020).

19 Mehta, A. R. et al. Mitochondrial bioenergetic deficits in C9orf72 amyotrophic lateral sclerosis motor neurons cause dysfunctional axonal homeostasis. Acta Neuropathol 141, 257–279, doi:10.1007/s00401-020-02252-5 (2021).

20 Leclech, C. & Villard, C. Cellular and Subcellular Contact Guidance on Microfabricated Substrates. Front Bioeng Biotechnol 8, 551505, doi:10.3389/fbioe.2020.551505 (2020).

21 Marcus, M. et al. Interactions of Neurons with Physical Environments. Advanced Healthcare Materials 6, 1700267, doi:10.1002/adhm.201700267 (2017).

22 Hagemann, C. et al. Automated and unbiased discrimination of ALS from control tissue at single cell resolution. Brain Pathol 31, e12937, doi:10.1111/bpa.12937 (2021).

23 Qin, D., Xia, Y. & Whitesides, G. M. Soft lithography for micro-and nanoscale patterning. Nature Protocols 5, 491–502, doi:10.1038/nprot.2009.234 (2010).

24 Artimovich, E., Jackson, R. K., Kilander, M. B. C., Lin, Y.-C. & Nestor, M. W. PeakCaller: an automated graphical interface for the quantification of intracellular calcium obtained by high-content screening. BMC Neurosci 18, doi:10.1186/s12868-017-0391-y (2017).

25 Gasperini, R. J. et al. How does calcium interact with the cytoskeleton to regulate growth cone motility during axon pathfinding? Molecular and Cellular Neuroscience 84, 29–35, doi:10.1016/j.mcn.2017.07.006 (2017).

26 Öztürk, Z., O’Kane, C. J. & Pérez-Moreno, J. J. Axonal Endoplasmic Reticulum Dynamics and Its Roles in Neurodegeneration %U https://www.frontiersin.org/article/10.3389/fnins.2020.00048. Front Neurosci 14, p%7 %8 2020-January-2029 %2029 Review %# %! Axonal ER dynamics and roles %* %<, doi:10.3389/fnins.2020.00048 %W %L (2020).

27 Rishal, I. et al. A Motor-Driven Mechanism for Cell-Length Sensing. Cell Rep 1, 608–616, doi:10.1016/j.celrep.2012.05.013 PMID - 22773964 (2012).

28 Harris, J., Julia Jolivet, R. & Attwell, D. Synaptic Energy Use and Supply. Neuron 75, 762–777, doi:10.1016/j.neuron.2012.08.019 (2012).

29 Pham, A. H., McCaffery, J. M. & Chan, D. C. Mouse lines with photo-activatable mitochondria to study mitochondrial dynamics. Genesis 50, 833–843, doi:papers3://publication/doi/10.1002/dvg.22050 (2012).

30 Spillane, M., Ketschek, A., Tanuja, Jeffery & Gallo, G. Mitochondria Coordinate Sites of Axon Branching through Localized Intra-axonal Protein Synthesis. Cell Rep 5, 1564–1575, doi:10.1016/j.celrep.2013.11.022 (2013).

31 Sheng, Z.-H. The Interplay of Axonal Energy Homeostasis and Mitochondrial Trafficking and Anchoring. Trends Cell Biol 27, 403–416, doi:10.1016/j.tcb.2017.01.005 (2017).

32 Campello, S. & Scorrano, L. Mitochondrial shape changes: orchestrating cell pathophysiology. EMBO reports 11, 678–684, doi:10.1038/embor.2010.115 (2010).

33 Westrate, L. M., Drocco, J. A., Martin, K. R., Hlavacek, W. S. & Mackeigan, J. P. Mitochondrial Morphological Features Are Associated with Fission and Fusion Events. PLoS ONE 9, e95265, doi:10.1371/journal.pone.0095265 (2014).

34 Santel, A. & Fuller, M. T. Control of mitochondrial morphology by a human mitofusin. J Cell Sci 114, 867–874, doi:10.1242/jcs.114.5.867 (2001).

35 Lewis, T. L., Kwon, S.-K., Lee, A., Shaw, R. & Polleux, F. MFF-dependent mitochondrial fission regulates presynaptic release and axon branching by limiting axonal mitochondria size. Nat Commun 9, doi:10.1038/s41467-018-07416-2 (2018).

36 Legati, A. et al. Current and New Next-Generation Sequencing Approaches to Study Mitochondrial DNA. J Mol Diagn, doi:10.1016/j.jmoldx.2021.03.002 (2021).

37 Kim, E. & Jung, H. Local mRNA translation in long-term maintenance of axon health and function. Curr Opin Neurobiol 63, 15–22, doi:10.1016/j.conb.2020.01.006 PMID - 32087477 (2020).

38 Gillingwater, T. H. & Wishart, T. M. Mechanisms underlying synaptic vulnerability and degeneration in neurodegenerative disease. Neuropathol Appl Neurobiol 39, 320–334, doi:papers3://publication/doi/10.1111/nan.12014 (2013).

39 Rossoll, W. & Bassell, G. J. Crosstalk of Local Translation and Mitochondria: Powering Plasticity in Axons and Dendrites. Neuron 101, 204–206, doi:10.1016/j.neuron.2018.12.027 (2019).

40 Schmidt, E. K., Clavarino, G., Ceppi, M. & Pierre, P. SUnSET, a nonradioactive method to monitor protein synthesis. Nature Methods 6, 275–277, doi:10.1038/nmeth.1314 (2009).

41 Jung, H., Yoon, B. C. & Holt, C. E. Axonal mRNA localization and local protein synthesis in nervous system assembly, maintenance and repair. Nat Rev Neurosci 13, 308–324, doi:papers3://publication/doi/10.1038/nrn3210 (2012).

42 Krichevsky, A. M. & Kosik, K. S. Neuronal RNA Granules. Neuron 32, 683–696, doi:10.1016/s0896-6273(01)00508-6 (2001).

43 Birsa, N. et al. FUS-ALS mutants alter FMRP phase separation equilibrium and impair protein translation. Science Advances 7, eabf8660, doi:10.1126/sciadv.abf8660 (2021).

44 Luisier, R. et al. Intron retention and nuclear loss of SFPQ are molecular hallmarks of ALS. Nat Commun 9, 2010, doi:papers3://publication/doi/10.1038/s41467-018-04373-8 (2018).

45 Tyzack, G. E. et al. Widespread FUS mislocalization is a molecular hallmark of amyotrophic lateral sclerosis. Brain 142, 2572–2580, doi:10.1093/brain/awz217 (2019).

46 Serio, A. et al. Astrocyte pathology and the absence of non-cell autonomy in an induced pluripotent stem cell model of TDP-43 proteinopathy. Proc Natl Acad Sci U S A 110, 4697–4702, doi:papers3://publication/doi/10.1073/pnas.1300398110 (2013).

47 Bilican, B., Serio, A., Shaw, C. E., Maniatis, T. & Chandran, S. Unpicking neurodegeneration in a dish with human pluripotent stem cells: one cell type at a time. Cell Cycle 12, 2339–2340, doi:papers3://publication/doi/10.4161/cc.25705 (2013).

48 Taylor, A. M. et al. A microfluidic culture platform for CNS axonal injury, regeneration and transport. Nature Methods 2, 599–605, doi:10.1038/nmeth777 (2005).

49 Neto, E. et al. Compartmentalized Microfluidic Platforms: The Unrivaled Breakthrough of In Vitro Tools for Neurobiological Research. The Journal of Neuroscience 36, 11573–11584, doi:10.1523/jneurosci.1748-16.2016 (2016).

50 Prokop, A. Cytoskeletal organization of axons in vertebrates and invertebrates. Journal of Cell Biology 219, doi:10.1083/jcb.201912081 (2020).

51 Shecket, G. & Lasek, R. J. Preparation of Neurofilament Protein from Guinea Pig Peripheral Nerve and Spinal Cord. Journal of Neurochemistry 35, 1335–1344, doi:10.1111/j.1471-4159.1980.tb09007.x (1980).

52 Vassilopoulos, S., Gibaud, S., Jimenez, A., Caillol, G. & Leterrier, C. Ultrastructure of the axonal periodic scaffold reveals a braid-like organization of actin rings. Nat Commun 10, doi:10.1038/s41467-019-13835-6 (2019).

53 Dubey, S. et al. The axonal actin-spectrin lattice acts as a tension buffering shock absorber. eLife 9, doi:10.7554/elife.51772 (2020).

54 Le Masson, G., Przedborski, S. & Abbott, L. F. A Computational Model of Motor Neuron Degeneration. Neuron 83, 990, doi:papers3://publication/doi/10.1016/j.neuron.2014.08.018 (2014).

55 Gopal, S. et al. Biointerfaces: Porous Silicon Nanoneedles Modulate Endocytosis to Deliver Biological Payloads (Adv. Mater. 12/2019). Advanced Materials 31, 1970086, doi:10.1002/adma.201970086 (2019).

56 Berg, S. et al. ilastik: interactive machine learning for (bio)image analysis. Nature Methods 16, 1226–1232, doi:10.1038/s41592-019-0582-9 (2019).

57 Carpenter, A. E. et al. Genome Biol. 7, R100, doi:10.1186/gb-2006-7-10-r100 (2006).

58 Goedhart, J. SuperPlotsOfData—a web app for the transparent display and quantitative comparison of continuous data from different conditions. Molecular Biology of the Cell 32, 470–474, doi:10.1091/mbc.e20-09-0583 (2021).

